# Zebrafish pigment cells develop directly from persistent highly multipotent progenitors

**DOI:** 10.1101/2021.06.17.448805

**Authors:** Masataka Nikaido, Tatiana Subkhankulova, Leonid A. Uroshlev, Artem J. Kasianov, Karen Camargo Sosa, Gemma Bavister, Xueyan Yang, Frederico S. L. M. Rodrigues, Thomas J. Carney, Hartmut Schwetlick, Jonathan H.P. Dawes, Andrea Rocco, Vsevelod Makeev, Robert N. Kelsh

## Abstract

Neural crest cells (NCCs) are highly multipotent stem cells. A long-standing controversy exists over the mechanism of NCC fate specification, specifically regarding the presence and potency of intermediate progenitors. The direct fate restriction (DFR) model, based on early *in vivo* clonal studies, hypothesised that intermediates are absent and that migrating cells maintain full multipotency^1–6^. However, most authors favour progressive fate restriction (PFR) models, with fully multipotent early NCCs (ENCCs) transitioning to partially-restricted intermediates before committing to individual fates^7–12^. Here, single cell transcriptional profiling of zebrafish pigment cell development leads to us proposing a Cyclical Fate Restriction mechanism of NCC development that reconciles the DFR and PFR models. Our clustering of single NCC Nanostring transcriptional profiles identifies only broadly multipotent intermediate states between ENCCs and differentiated melanocytes and iridophores. Leukocyte tyrosine kinase (Ltk) marks the multipotent progenitor and iridophores, consistent with biphasic *ltk* expression^13–15^. Ltk inhibitor and constitutive activation studies support expression at an early multipotent stage, whilst lineage-tracing of *ltk-*expressing cells reveals their multipotency extends beyond pigment cell-types to neural fates. We conclude that pigment cell development does not involve a conventional PFR mechanism, but instead occurs directly and more dynamically from a broadly multipotent intermediate state.

## Main

NCCs generate an astonishing diversity of cell-types, including cranial skeletogenic fates, peripheral neurons and glia and, except in mammals, multiple pigment cells ^16–19^. Fish pigment cells include black melanocytes (M), yellow xanthophores (X), and reflective iridophores (I)^20^. In zebrafish, pigment cell (*M*elanocyte, *I*ridophore, *X*anthophore) fate specification is an important test case, since a PFR model with distinct multipotent (MIX) and bipotent (melanoiridoblast, MI) intermediates has been suggested^13–15,21–23^. A long-standing, yet untested, hypothesis proposes that all vertebrate pigment cell-types share a common, and exclusive, progenitor, the chromatoblast (MIX)^21^. Analysis of mutants for the *microphthalmia-related transcription factor a (mitfa)* gene, encoding a master regulator for melanocytes^24,25^, were consistent with a bipotential MI progenitor in zebrafish^23^. These and complementary studies in medaka^26–28^ lead to a widely-accepted, but untested, PFR model of pigment cell development (Fig. 1). However in mouse and chick, shared progenitors for melanocytes and glia have been proposed^29,30^, and whilst in mouse migrating NC cells retain multipotency^31^, neural derivatives appear to originate via a PFR mechanism^32^.

**Fig. 1:**
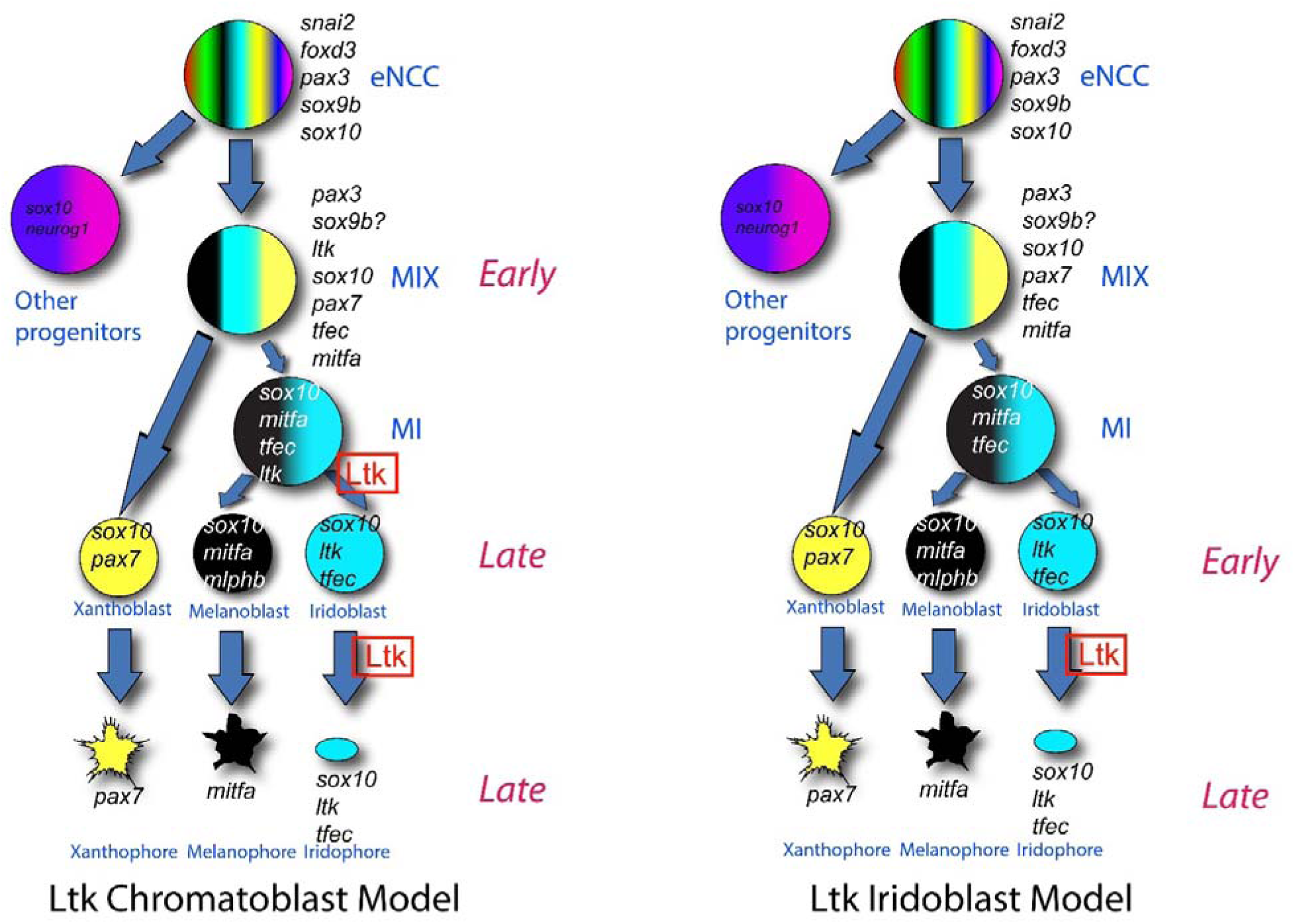
PFR models for zebrafish pigment cell development from neural crest. Models show eNCCs generating multipotent (MIX) and bipotent (MI) intermediates en route to generating melanocytes and iridophores, and distinguish timing of expression and role of Ltk signaling: **a**, Ltk Chromatoblast Model **b**, Ltk Iridoblast Model. In these schema, potency of cells (number of fill colours) at different stages in PFR of the pigment cell lineages decreases down the diagram (i.e. with time), reflecting PFR. Expression of *ltk* is indicated by italics (*ltk*); other key marker genes are indicated too. Ltk function (signaling activity) is indicated in Roman script (Ltk, boxed). Thus, in the Ltk Chromatoblast Model, *ltk* is expressed in all chromatoblasts (MIX) and melanoiridoblasts (MI)(‘Early’), but Ltk signaling is activated only in a subset of them, driving iridophore lineage specification (iridoblast specification). Continuing *ltk* expression in iridoblasts and iridophores (‘Late’) has *late* role in iridophore differentiation, proliferation and/or survival. In the Ltk Iridoblast Model, early phase expression represents iridoblasts (‘Early’), where it functions in differentiation or survival. Late expression reflects ongoing expression and ongoing function in differentiated iridophores. Experimental assessment using Ltk inhibitor treatment reveals early role in iridophore fate specification and later role in differentiated cells, supporting Ltk Chromatoblast Model (**a**) (Extended Data Fig. 9).

### Single cell transcriptional profiling of zebrafish NCCs fails to identify putative partially-restricted intermediates

We investigated the transcriptional profiles of 1317 zebrafish NCCs throughout pigment cell development (18-72 hpf). To obtain sensitive quantitation of key mRNA expression, we used the NanoString platform and a set of 45 genes, focused on those known or suspected to have important roles in melanocyte or iridophore fate choice, but including multiple markers of early NCCs, plus key markers of other major NC-derived fates (Supplementary Table 1). We profiled FACS sorted cells from *Tg(Sox10:Cre)^ba74^xTg(hsp70l:loxP-dsRed-loxP –Lyn-Egfp^tud107Tg^* embryos at 8 time-points in which NCCs and their derivatives are labelled with eGFP after a brief heat shock (Figure 2a). Our sample consisted of WT cells isolated at time points between 18-72 hpf, plus four distinct sets of control cells: FACS-sorted NCCs from dissected tails of WT embryos at 24 hpf (expected under default PFR model to be enriched for early NCCs and MIX); NCCs from *sox10* mutants at 30 hpf (enriched for cells trapped in MIX state^13,15,33^); and gradient centrifugation-purified differentiated melanocytes and iridophores from 72 hpf WT embryos. The results of NanoString profiling went through a series of quality control tests and rounds of batch correction and normalization, resulting in 731 cells profiled for expression of 42 genes. The set included 444 experimental (denoted “regular”, reg) WT cells, and 287 control cells (108 WT tail, 135 *sox10* mutant, 25 iridophores (contI) and 19 melanocytes (contM)). To identify cells with similar expression profiles we conducted cell clustering by gene expression profiles using shared nearest neighbour clustering algorithm implemented in the Seurat R package. We employed control cell types to obtain a natural measure of gene expression variation within a cell type and selected parameters of the clustering algorithm so that the “control clusters” containing most of the control melanocytes and iridophores respectively, had the largest proportion of the control cells in the respective cluster. Initial clustering generated 10 clusters, but some pairs of clusters had similar expression profiles; we used the Seurat`:ValidateClusters` procedure to merge similar clusters, resulting in 7 clusters (Figure 2b,c). Identification of specific clusters 1-3, 5 and 6 was facilitated by a combination of enrichment for control cell-types (Extended Data Fig. 1) and expression of key markers (Extended Data Fig. 2,3), and these are named accordingly. Thus, one cluster contained all control melanocytes, as well as 18 other cells, while another contained 20 (80%) of the control iridophores, plus 29 other cells (Extended Data Fig. 1a). These clusters respectively expressed melanocyte (e.g. *mlphb*, *oca2*, *slc24a5*, *tyrp1b*, and *pmel*) and iridophore (e.g. *pnp4a* and *ltk*) gene markers (Extended Data Fig. 1d,3,4). As expected, the expression profiles of control cells (contI and contM) and the other cells (regI and regM) in each of these two clusters were near identical (Extended Data Fig. 1c). Interestingly, melanocytes were clearly segregated into two sub-clusters, each containing similar numbers of control melanocytes; both express characteristic melanocyte differentiation genes, but one (mostly consisting of control melanocytes) additionally expresses early NCC markers (*snail2, sox9b*) and genes functionally associated with iridophore and xanthophore development (*ltk, tfec, pax7a*), plus *foxo1a* (Extended Data Fig. 4). One further cluster is tentatively identified as xanthophores by the combination of high level expression of *pax7a* and *pax7b* (Fig. 2b,c; Extended Data Fig. 2,3) ^34,35^. The three pigment cell clusters were very stable and never merged with other clusters up to very high threshold values (0.92) of the Seurat:`ValidateClusters` procedure. As expected, given the targeted focus on melanocytes and iridophores in our Nanostring profile gene selection, clusters corresponding to neural and skeletogenic (cartilage) fates were not identifiable, but likely such cells are included in the two large unassigned clusters 4 and 7, which consequently are likely much more heterogeneous than is apparent with our marker set (Fig. 2b; Extended Data Fig. 2,3).

**Fig. 2:**
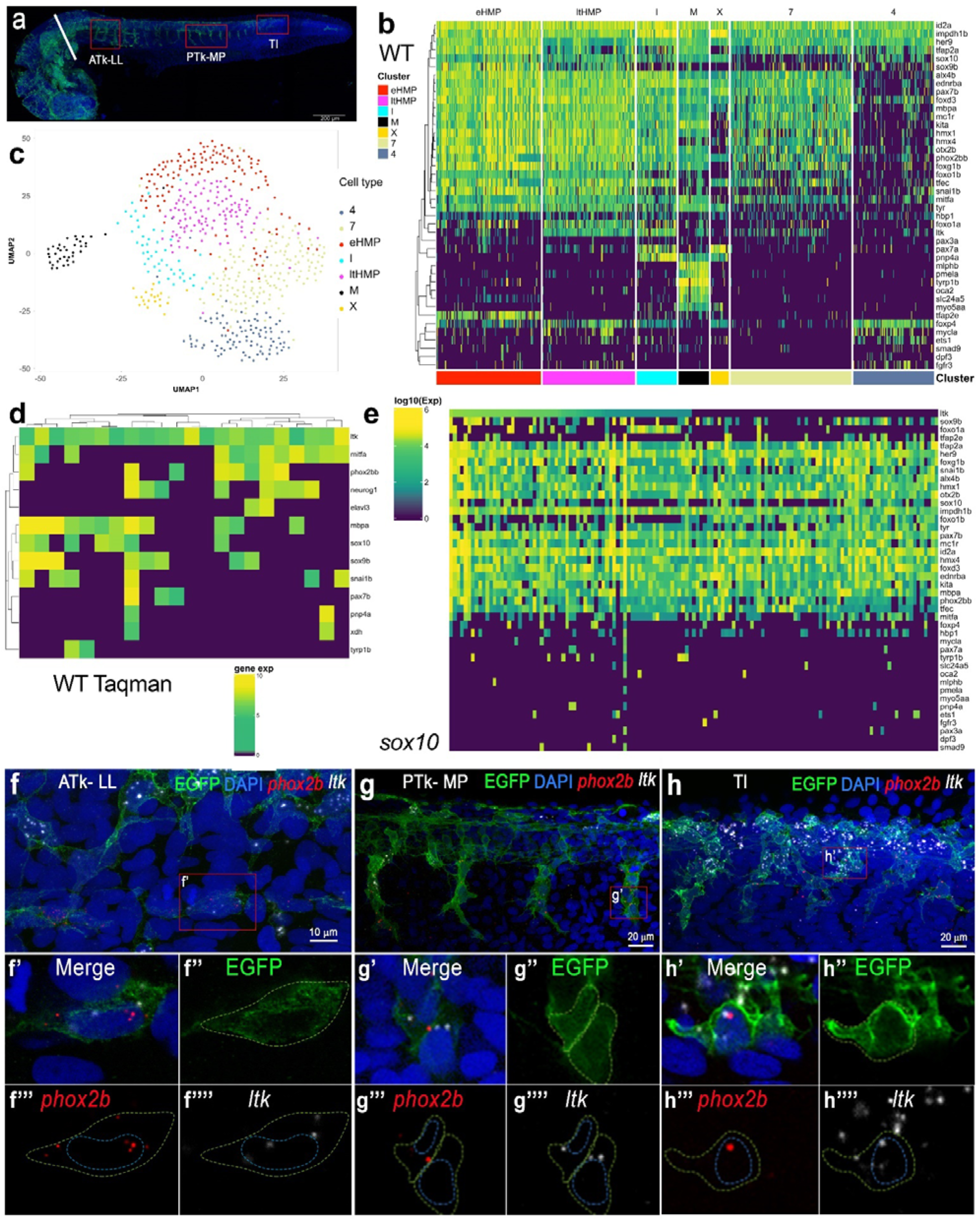
Highly multipotent, but not partially fate-restricted, intermediates are readily detected in zebrafish embryos. **a**, Whole-mount immunodetection of EGFP in *Tg(Sox10:Cre)^ba74^xTg(hsp70l:loxP-dsRed-loxP –Lyn-Egfp^tud107Tg^* to show source of single cells for NanoString profiling (24 hpf stage shown here); embryos at 30 hpf or older had heads removed by severing behind otic vesicle (white bar). Red boxes identify regions imaged in close-up in panels **f-h** to show anterior trunk lateral migration pathway and posterior lateral line nerve (ATk-LL), posterior trunk medial pathway (PTk-MP) and tail (Tl). **b-e**, Single cell profiling of zebrafish NCCs reveals unexpected absence of partially-restricted pigment cell progenitors. **b**, Heat maps showing NanoString profiles of clusters identified in **c**; see also Extended Data Fig. 3 for violin-plot representation. **c**, NCC profile clustering, clustered and visualized in 2D by UMAP. Clusters are identified by distinctive expression profiles: eHMP (red), early highly multipotent progenitors; ltHMP (magenta), late highly multipotent progenitors; I (cyan), iridophores; X (gold), xanthophores; M (black), melanocytes; clusters marked with numbers not identified due to lack of distinctive marker gene profiles. **d**, Heat map showing independent qRT-PCR (Taqman assay) assessment of overlapping fate specification gene expression in 24 hpf WT embryos. Only cells with detectable *ltk* expression shown; for full profiles of all cells, see Extended Data Fig. 6. Note that individual cells usually combine expression of *mitfa, pax7b, tfec, ltk, sox10, phox2bb*, with multiple genes being detected in each individual cell. **e**, Heat map of *sox10* mutant cell NanoString transcriptional profiles, ordered by *ltk* expression levels. **f-h**, RNAscope expression analysis reveals co-expression of pigment cell and neuronal fate specification genes *in vivo* in 27 hpf *Tg(Sox10:Cre)^ba74^xTg(hsp70l:loxP-dsRed-loxP–Lyn-Egfp^tud107Tg^* embryos after brief heat-shock to express EGFP. **f**, **g**, **h**, RNAscope FISH detection of *phox2bb* (red) and *ltk* (white) is shown in confocal lateral view projections of the lateral line (LL) in the anterior trunk (ATk; **f**), medial pathway (MP) in the posterior trunk (PTk; **g**) and tail (Tl; **h**). Insets (f’-h’) show co-expression of *phox2bb* and *ltk* in EGFP-labelled cells. Merge and individual channels for a single focal plane for each inset is shown in f’-h’’’’. Yellow dashed line shows membrane cell border. Blue dashed line shows outline of nucleus for NCCs as revealed by DAPI (blue in merge). Scale bar dimensions are as indicated.

Due to the developmental gradient along the anteroposterior axis, NCCs isolated from tails at 24 hpf are enriched for developmentally early cell-types. One cluster was highly enriched with tail cells (Extended Data Fig. 1a), and showed high level expression of two known early NCC markers (*snail2, sox9b*)^36–38^, but also *tfap2a* and *tfap2e*, both required early in melanocyte development^39^(Fig. 2b; Extended Data Fig. 1d,2,3). Consequently, we interpreted this cluster as the earliest stage isolated by our *sox10-*dependent labelling technique (note that *sox10* expression begins after *sox9b* and *snai2*, which are both downregulated in most *sox10*-expressing cells^15^), naming them early highly multipotent progenitors (eHMP). We used *slingshot* ^40^ to build pseudotime trajectories of pigment cell differentiation, starting from the eHMP (Extended Data Fig. 5). Trajectories for both melanocyte and iridophore differentiation went through another cluster which also contained some tail cells and showed a similar expression profile; we named these cells late highly multipotent progenitors (ltHMP). The ltHMP cells were distinguished from eHMPs by increased expression of *ltk* and lower levels of *snai2* and *sox9b* (Fig. 2b; Extended Data Fig. 1d,2,3), consistent with the known progression of marker expression in zebrafish NCCs^13–15^, and also by a striking decrease of *tfap2e*. In contrast, both eHMP and ltHMP show prominent expression of key fate specification genes (*mitfa, pax7b, tfec, phox2bb, sox10*) for diverse pigment cell and neural fates. Both eHMP and ltHMP cells are identified from all stages, even up to 72 hpf, suggesting that they are not just found in premigratory NCCs, but persist at later stages. Interestingly, when *sox10* mutant cells are included, we observed that they were not separated from WT cells, but occupy part of the central cloud (Extended Data Fig. 1b) and are effectively excluded from the differentiated pigment cell clusters. As expected given the strong failure of pigment cell fate specification in *sox10* mutants^13–15,17,33^, mutant cells generally lack pigment cell markers (e.g. *pnp4a*, *pax7a, mlphb*, *oca2*, *pmel*, *slc24a5*, and *tyrp1b*) and show somewhat reduced expression of some fate specification (e.g. *mitfa*) and early markers (e.g. *sox9b*). They show retained expression of other early markers (e.g. *snai2*) and broadly-expressed genes (e.g. *id2a, impdh1b, ednrba, alx4b*)(Fig. 2e; Extended Data Fig. 1d,2). In general, *sox10* mutant cells are more homogeneous than WT cells and form a single cluster if optimal parameters of cluster validation for WT clustering are used. Broadly, their expression profile resembles the HMP clusters seen in WTs. A subset of *sox10* mutant cells show elevated levels of *ltk* expression (Fig. 2e), comparable to the ltHMP cluster (Fig. 2b), and in agreement with semi-quantitative observations by whole-mount in situ hybridization of cells trapped in a premigratory position^15,17^. Together, these observations are consistent with the previous deduction that mutant cells with the potential to adopt pigment cell fates (indicated by expression of *tfec*, *ltk*, *pax7b,* and *tyr*; Fig. 2e; Extended Data Fig. 1d,2) are prevented from proceeding along the pigment cell differentiation pathways, and remain in a multipotent progenitor state characterised by expression of *ltk*, *transcription factor EC (tfec)* and *sry-related HMG box10 (sox10)*^13–15,33^. However, most cells of this state also express *phox2bb* (Fig. 2e; Extended Data Fig. 1d,2), indicating a previously unanticipated potential to adopt at least some neuronal fates as well, although this potential is not realized *in vivo* because it is blocked by the absence of functional Sox10^41^.

We consider that gene expression profile provides a molecular correlate of (minimal) cell-potency, with expression of known fate specification genes indicating potential^42^. It is striking that our analysis failed to identify clusters of cells corresponding to either the chromatoblast or melanoiridoblast predicted under the PFR model (Fig. 2b,c; Extended Data Fig. 2,3). Thus whilst both eHMP and ltHMP clusters express genes with known role in fate specification of each pigment (*mitfa, pax7b, tfec, ltk)*, neuronal (*phox2bb)* and glial (*sox10*) cell-type, no clusters show specific combinations consistent with MIX (*mitfa, pax7b, tfec, ltk,* but not *phox2bb*) nor MI (*mitfa, tfec, ltk,* but not *pax7b,* or *phox2bb*) intermediates. As an independent validation of the overlap of *ltk* and other pigment fate specification genes with early neuronal fate specification genes, we used TaqMan qRT-PCR to evaluate a further sample of more than 100 single NCCs, FACS-sorted from 30 hpf WT embryos, examining both *phox2bb* (sympathetic and enteric neuron fate specification) and *neurog1* (sensory neuron) expression (Fig. 2d; Extended Data Fig. 6). Out of 21 cells expressing detectable *ltk* (with Ct≤29), 12 (57%) showed detectable *mitfa*, whilst 12 (57%) showed one or both of *phox2bb* and *neurog1*; if we include the Schwann cell marker *mbpa* then 17 (81%) of *ltk*-expressing cells show one or more of these neural fate markers (Fig. 2d). Examining the full data-set, we note how 79% and 85% of cells expressing *phox2bb* or *neurog1* respectively express at least one of the key pigment cell fate specification genes *ltk, mitfa* or *pax7b* (Extended Data Fig. 6). Together our data suggests the surprising possibility that broad multipotency is retained in most progenitor cells.

### Co-expression of key fate specification genes in situ

To further validate these unexpected findings, we used our optimised RNAscope in situ hybridization protocol to examine co-expression of pigment cell and neuronal fate specification genes. We focused on testing *phox2bb* co-expression patterns that were unexpected under the PFR model. Expression of *phox2bb* in zebrafish has been previously reported as restricted to NCCs migrating along the developing gut from c 24 hpf onwards, interpreted as progenitors of the enteric ganglia, and sympathetic ganglial progenitors in the ventral medial pathway from 36 hpf^43–45^. Analysis of *phox2bb* morphants suggests it is required for fate specification of enteric and sympathetic neurons^41,45^. Using RNAscope, we detect *phox2bb* expression in 27 hpf zebrafish within many premigratory NCCs, but also some migrating NCCs on the medial pathway, and even in many NCCs associated with the posterior lateral line nerve (Fig. 2f-h; Extended Data Fig. 7). Furthermore, in all of these locations, cells also expressing *ltk*, a key iridophore specification gene, are readily found ^15^(Fig. 2f-h; Extended Data Fig. 7). Thus, *phox2bb* expression can be detected at low levels in cells from premigratory stages, long before formation of sympathetic or enteric ganglial progenitors, but consistent with the expected autonomic neuron potential of premigratory NCCs (Fig. 2h; Extended Data Fig. 7i). More strikingly, that autonomic neuron potential is then maintained in a widespread manner, including both ventralmost cells on the medial migration pathway (Fig. 2g; Extended Data Fig. 7g,h) and in putative Schwann cell progenitors associated with the lateral line nerve (Fig. 2f; Extended Data Fig. 7f). The expression in putative Schwann Cell Precursors of the posterior lateral line nerve is intriguing, suggesting retained autonomic neuron potential that is unlikely to be realized *in vivo*.

### Biphasic expression of *ltk* distinguishes multipotent iridophore progenitors from iridoblasts

Expression of *leukocyte tyrosine kinase* (*ltk*) was particularly striking in our single cell profiles, showing relatively high expression in both ltHMPs and iridophores (Fig. 2b). In the context of previous work, this suggested a series of further tests of the model emerging from our single cell expression profiling, as now described. *ltk* encodes a receptor tyrosine kinase, and analysis of loss- and gain-of-function mutations indicated that Ltk signalling drives iridophore fate specification^15,46,47^. We had postulated two phases of expression of *ltk* in NCCs: Late phase expression (from around 26 hours post-fertilization (hpf) in the posterior trunk, later In tail) corresponds to the differentiating iridophore lineage, while early phase expression (around 22-24 hpf in the trunk) in premigratory NCCs represents multipotent progenitors (Fig. 1a) ^13–15^. An alternative model postulates that both early and late phase *ltk* expression is in cells committed to the iridophore lineage (Fig. 1b). Interestingly, in our Nanostring profiles *ltk* shows covariance with two clusters of genes (iridophore markers (e.g. *pnp4a, impdh1b*) and broadly expressed genes; Extended Data Fig. 8), suggesting biphasic expression, consistent with this proposal. To test directly the biphasic expression model, we used a small molecule inhibitor of Ltk^46^ to disrupt Ltk signalling in zebrafish embryos at different stages; allowing for the spatiotemporal differences in NCC development along the anteroposterior axis, we find distinct effects of inhibition of early and late phase expression, with the former restricting the number of iridophores specified, and the latter controlling the expansion of these cells, likely by proliferation to form clones (Extended Data Fig. 9).

### Constitutive Ltk signalling drives iridophore fate at expense of other pigment cell fates

As a further test, we predicted that, if Ltk activity drives iridophore fate from multipotent progenitors, expression of constitutively active Ltk signalling in NCCs would promote ectopic and supernumerary iridophore formation at the expense of other fates. We used constitutive activation of Ltk through generation of an N-terminal fusion with human Nucleophosmin, NPM-Ltk (Fig. 3a), which drives NCCs to an iridophore fate when expressed using the zebrafish *sox10* promoter ^46^. We generated a plasmid, *Tg(Sox10:NPM-ltk, egfp)* (hereafter, *Tg(sox10*:*NPM*-*ltk*)) which uses the *sox10* promoter to drive expression of both NPM-Ltk and the lineage tracer, enhanced GFP (Fig. 3a). As a negative control, we created a dead kinase (DK) version of *sox10*:*NPM*-*ltk* by exchanging lysine 943 in the kinase domain of wild type zebrafish Ltk protein with arginine (K943R, Fig. 3a)(plasmid *Tg(Sox10:NPM-ltk_K943R, egfp)*, here referred to as *Tg(sox10*:*NPM*-*ltk(DK))*) based on the previously reported activity in human *ALK* ^48^. To assay the fate of constitutively expressing cells, we focused on whether the *Tg(sox10*:*NPM*-*ltk*) construct might redirect cells from a melanocyte to an iridophore fate, because the melanocyte fate is readily characterized by colour, and because the reciprocal fate switch is hypothesised to underlie the increased number of iridophores documented in *mitfa*/*nacre* mutants^22^. Whilst *Tg(sox10*:*NPM*-*ltk*) readily induced precocious (Fig. 3c) and ectopic (Fig. 3c’,c’’) iridophores at this stage, these were almost undetected at 60 hpf in embryos injected with *Tg(sox10*:*NPM*-*ltk(DK))* (Fig. 3b, 3b’), showing that *Tg(sox10*:*NPM*-*ltk(DK))* has minimal NPM-Ltk activity (Extended Data Table 1). We injected embryos with *Tg(sox10*:*NPM*-*ltk*) or with *Tg(sox10*:*NPM*-*ltk(DK))* and scored embryos having normally elongated body axis at 60 hpf for the presence or absence of additional iridophores, and for GFP-positive melanocytes (Extended Data Table 1; Fig. 3d-3f’). As expected, in control embryos injected with DK version, we did not find embryos with additional iridophores. In the 50 *Tg(sox10*:*NPM*-*ltk(DK))* injected embryos that were GFP-positive, we identified 11 GFP-positive melanocytes (Table 1, Fig. 3e, 3e’). In striking contrast, we found only one GFP-positive melanocyte in the 53 GFP-positive embryos injected with *Tg(sox10*:*NPM*-*ltk*), a significant reduction (Table 1; Chi-squared test, P=0.00233). We conclude that active Ltk signalling in NCCs is largely incompatible with melanocyte development, instead driving cells to adopt an iridophore fate. Consistent with this interpretation, the single melanocyte obtained in an embryo injected with *Tg(sox10*:*NPM*-*ltk*) was poorly differentiated, being small, round and less dendritic than those in control embryos, and was found in an embryo that had no additional iridophores, suggesting weaker activation of Ltk signal therein. These data are consistent with the idea that Ltk signalling in a multipotent progenitor cell drives iridophore fate choice at the expense of other fates.

**Fig. 3:**
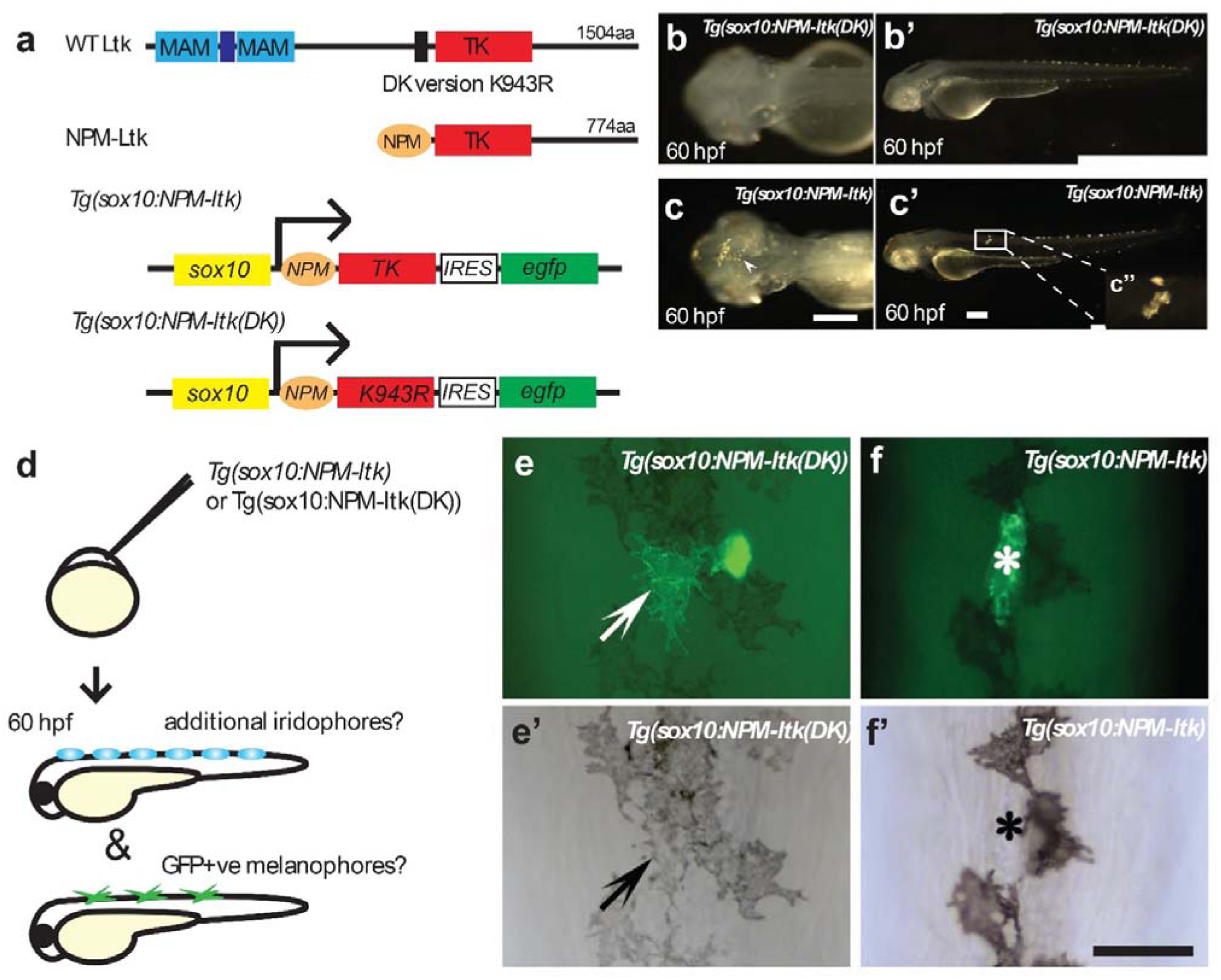
Active Ltk signaling is incompatible with melanophore development. **a**, Schematic drawings of wild-type Ltk and NPM-Ltk fusion protein, and of NC expression construct *Tg(sox10*:*NPM*-*ltk*); see text for details of these and of negative control kinase dead construct, *Tg(sox10*:*NPM*-*ltk(DK))*. **b-c’**, Validation of constitutively-activated Ltk and dead kinase control. Embryos injected with *sox10*:*NPM-ltk DK* (**b, b’**) and *Tg(sox10*:*NPM*-*ltk*) (**c, c’**) at 60 hpf. Precocious (arrowhead in **c**) and ectopic (inset **c”**) iridophores are shown. Melanisation was inhibited by PTU treatment to enhance detection of iridophores. **b, c** Dorsal views with anterior to the left. **b’, c’** left side views with dorsal to the top. Scale bars: 100 µm. **d-f’**, Ltk activity is inconsistent with melanocyte differentiation. **d**, Schematic drawing of experimental procedure. DNA constructs were injected into embryos at 1-cell stage. Embryos were cultured until 60 hpf, scored for precocious/ectopic iridophore formation, and subjected to anti-GFP antibody staining. **e-f’**, GFP-positive melanophore in control embryos injected with *Tg(sox10*:*NPM*-*ltk(DK))* (arrows in **e, e’**); a second GFP-positive cell was an iridophore based upon its position and shape. In embryos injected with *Tg(sox10*:*NPM*-*ltk*), GFP-positive cells were almost invariably not melanised (asterisks in **f, f’**). All views show dorsal midline, anterior to the top. Panels **e** and **f** are fluorescent images merged with bright field images and **e’** and **f’** are bright field images for **e** and **f**, respectively. Scale bar: 50 µm.

**Table 1.** Activated Ltk signalling suppresses melanophore formation.

|  | No. of observed<br>embryos | No. of GFP-positive<br>embryos in observed<br>embryos | No. of GFP-positive<br>melanophores in<br>GFP-positive<br>embryos<sup>(1)</sup> |
| --- | --- | --- | --- |
| *Tg(sox10:NPM-*<br>*ltk(DK))* | 215 (100%) | 50 (23%) | 11 |
| *Tg(sox10:NPM-ltk)*<br>(+irido)<sup>(2)</sup> | 95 (100%) | 37 (39%) | 0 |
| *Tg(sox10:NPM-ltk)*<br>(-irido)<sup>(2)</sup> | 142 (100%) | 16 (11%) | 1 |

Notes: <sup>(1)</sup> Embryos were raised in a low dose of PTU (25% of normal dose, i.e. 0.00075%) to partially inhibit melanin synthesis, so that GFP fluorescence in the living cells of injected embryos could be readily assessed in melanophores.
(2) “+/-irido” here means presence or absence of precocious/ectopic iridophores in injected embryos, respectively.

### *ltk*-expressing cells generate non-ectomesenchymal NC derivatives

To test directly the proposed multipotency of *ltk*-expressing cells, we used fate-mapping using transient expression of an *ltk:gfp* transgene, examining the prediction that green fluorescent protein (GFP) expression under the *ltk* transcriptional regulatory regions should label all pigment cells, but also other (e.g. neural) derivatives as well. We used BAC recombineering techniques^49,50^ to make a reporter construct for the *ltk* gene, *TgBAC(ltk:gfp)* (Fig. 4a). As a positive control construct to test we could label each NCC derivative when GFP was expressed in early NCCs, we used a *sox10*:*gfp* PAC (P1 phage-derived artificial chromosome; see Materials and Methods for details)(Fig. 4a). We injected each of these two constructs into *Tg(neurogenin1(8.3):rfp* transgenic fish, in which red fluorescent protein (RFP) is expressed in developing dorsal root ganglion (DRG) neurons in embryos from as early as 2 dpf^51^ to facilitate identification of GFP reporter expression in early stage DRG neurons, since our previous studies showed that *sox10-*driven GFP perdurance in DRG does not usually last beyond c. 48 hpf^52^. The presence of GFP expression in all NCC-derived cartilage, pigment cell and neural cell-types was assessed (see Materials and Methods for details of criteria). The *TgPAC(sox10:gfp)* construct labelled all expected neural crest derivatives as found by single cell labelling studies (Table 2; Fig. 4c)^18,19,33^. The results for embryos injected with the *TgBAC(ltk:gfp)* construct were striking (Table 2; Fig. 4b). Whilst many GFP-positive iridophores were seen in embryos injected with the *TgBAC(ltk:gfp)* construct, GFP expression was clearly not restricted to this cell-type, consistent with our expectation that early phase *ltk* expression represented a multipotent progenitor. Strikingly, we saw expression of GFP in numerous other pigment cells, including both melanocytes and xanthophores, but also in glial and neuronal derivatives, specifically DRG sensory neurons and enteric neurons. This observation indicated other NCC-derivatives, such as glial and neuronal cells, are also derived from early *ltk*-positive cells.

**Fig. 4:**
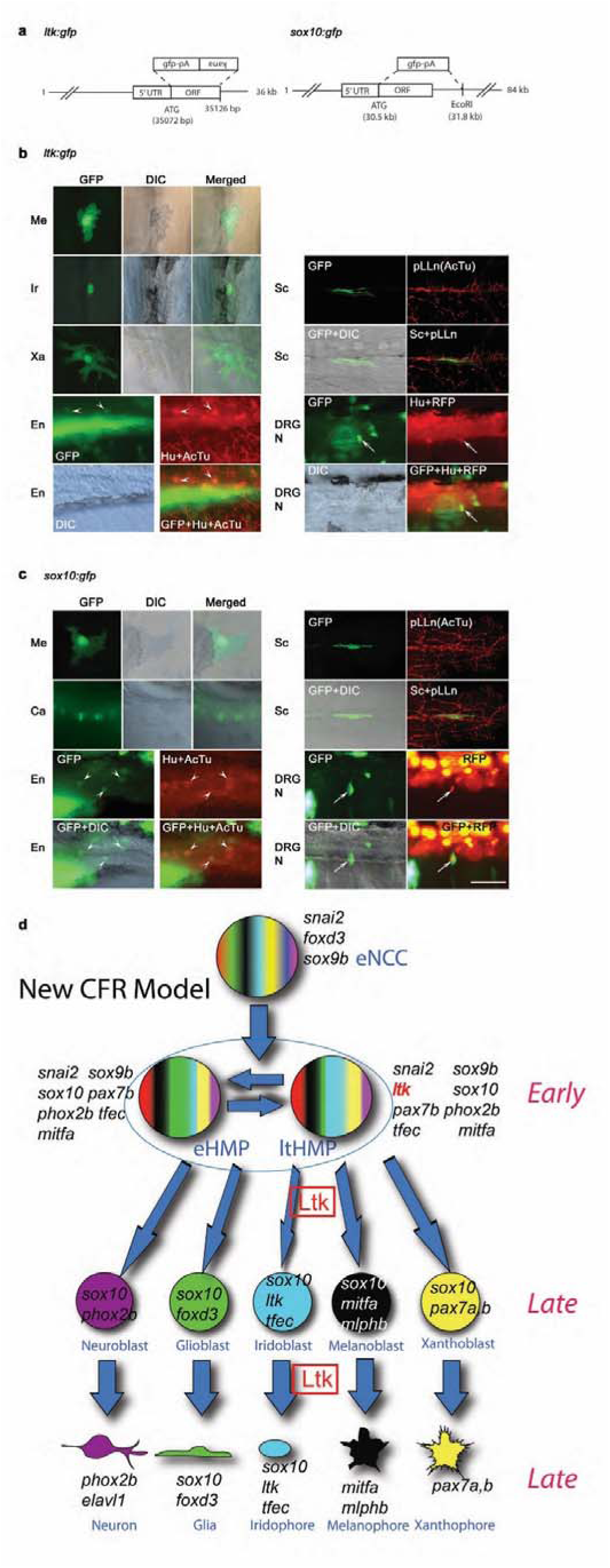
Genetic fate mapping using a *TgBAC(ltk:gfp)* construct identifies pigment cells, but also Schwann cells and neurons, as derived from *ltk-*expressing cells. **a**, Schematic drawings of reporters. In both constructs, *gfp* cDNA with polyA site (*pA*) is inserted at the site of the first methionine. See Materials and Methods for details. **b**, Cells labeled by *TgBAC(ltk:gfp)*, shown at 2 dpf (DRG neuron), 3 dpf (melanocyte) or 4 dpf (others). All pigment cells are labeled by GFP or anti-GFP antibody, and observed by bright field (bright) and immunofluorescence (GFP) microscopy; combined bright field and fluorescent images also shown (merged). Schwann cell around the posterior lateral line nerve is also detected by anti-GFP antibody (GFP), and pLLn, is detected by anti-acetylated tubulin antibody (AcTu). Enteric and DRG sensory neurons are labelled by GFP, with enteric neurons identified by position (revealed by Differential Interference Contrast (DIC)) and anti-Elavl1 staining (Hu)(arrowheads; anti-AcTu also detected), whilst DRGs are identified by position and anti-Elavl1 staining (Hu) and RFP driven by *neurog1* promoter (arrows) (see Materials and Methods). DRG N, DRG neuron; En, enteric neuron; Ir, Iridophore; Me, melanocyte; pLLn, posterior lateral line nerve; Sc, Schwann cell; Xa, Xanthophore. **c**, Cells labeled by *TgPAC(sox10*:*gfp)*. In addition to pigment cells and Schwann cells, lower jaw cartilage (Ca), enteric neurons and DRG are also labeled. For quantitation, see Table 2. Scale bar: 50 µm. **d**, New Cyclical Fate Restriction Model of pigment cell development. Pigment cells derive from Highly Multipotent Progenitor (HMP) cells, which are envisaged as cycling through multiple sub-states; for simplicity, only two of these (eHMP and ltHMP) are distinguished here, based principally upon the level of expression of *ltk* (see Fig. 2b). Early and late phase Ltk expression reflects that in the ltHMP and iridoblast/iridophore respectively, with Ltk function (Ltk, boxed) required early for specification of iridophore lineage from ltHMP and late for ongoing differentiation/proliferation. See Fig. 1 legend for key.

**Table 2.**
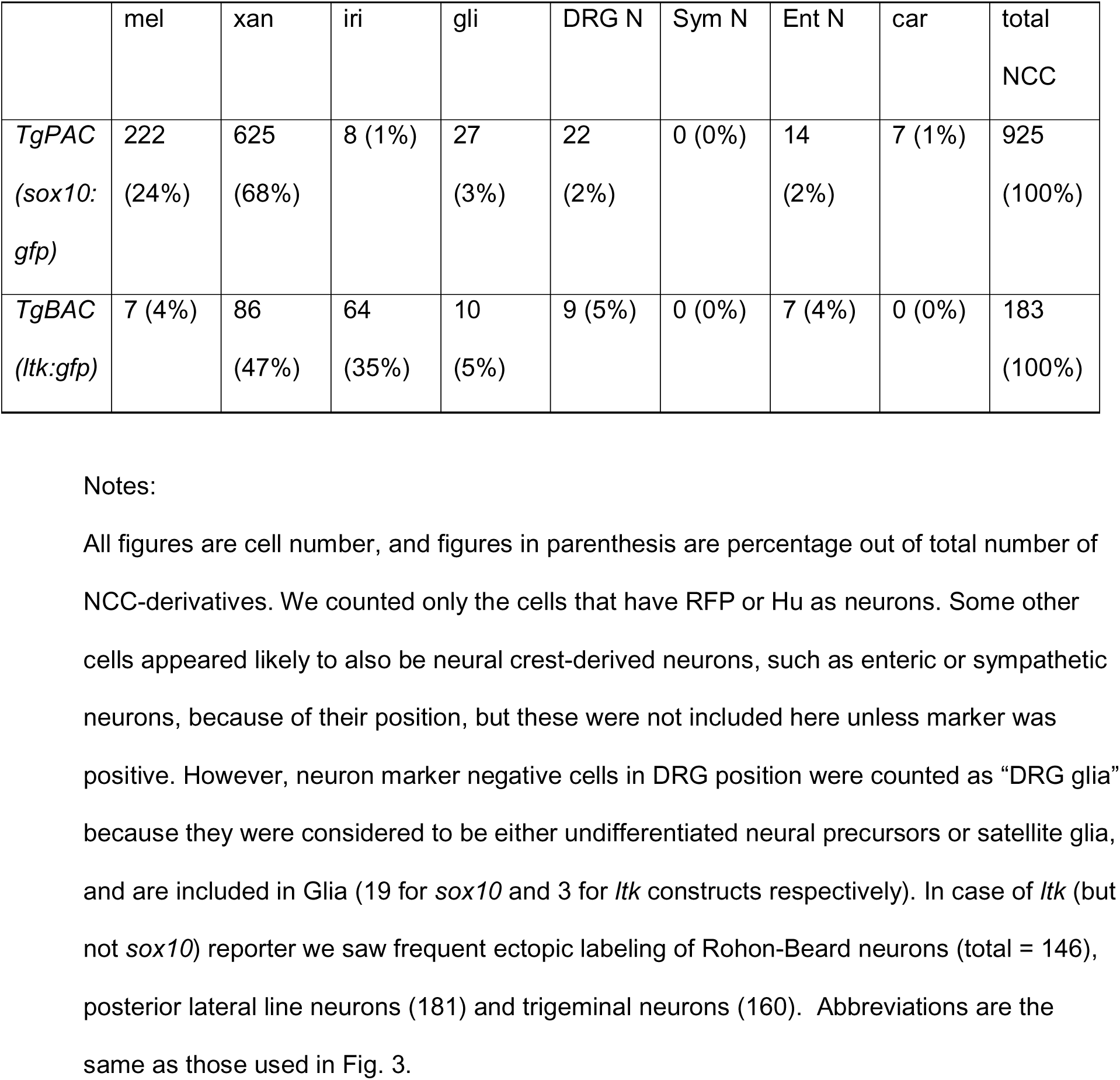
*ltk-*expressing cells generate neural and all pigment cell-types.

## Discussion

How NCCs adopt individual fates from their extensive repertoire has remained an enigma. Recent adoption of a PFR model has been most well-worked out for the adoption of pigment cell fates in zebrafish, where two distinct key intermediates were proposed (Fig. 1). Our data provides direct assessment of aspects of this model, with the unexpected conclusion that the identified progenitors (eHMP and ltHMP) showing a transcriptional profile including all key fate specification genes assessed, strongly indicating broader multipotency than expected under the PFR model; conversely, the expected multiple intermediates (MIX and MI) were not detectable *in vivo*. We and others have shown the crucial role for Ltk signalling in fate specification of iridophores^15,46,47^. Here we demonstrate experimentally that early phase expression of *ltk* in premigratory NCCs is necessary for this fate choice decision, and that Ltk signalling can suppress other fates, at least for the melanocyte fate assessed here. Lineage-tracing of the fate of these early phase *ltk* expressing cells demonstrates that early phase *ltk* expression does indeed mark multipotent progenitors, which generate glia and all types of pigment cells, but also at least some neuronal cells. An alternative explanation, that the *ltk* expressing NCCs are a mixture of cells of distinct potencies (i.e. broad multipotency is true only of the population, but not of individual cells) is inconsistent with our single cell profiling data, which readily identified a large cluster of cells showing overlapping expression of key genes underpinning diverse fate choices, including neural fates, reflecting their broad multipotency. A corollary of this observation is the insight that fate specification of an individual cell-type depends as much on repression of inappropriate fate-specific transcription factors (e.g. *phox2bb* in a melanocyte), as on upregulation of appropriate ones (e.g. *mitfa*)^13,14,42^.

We have been unable to identify clusters that would correspond to either the MIX or MI intermediates previously hypothesized. An alternative explanation is that our limited gene set lacked the key markers, although this seems unlikely since the markers used were mostly selected for their known crucial roles in fate specification of neural crest-derived cell-types. Nevertheless, further testing of our hypothesis will come from comprehensive (i.e. deep) single cell RNA-seq profiling of zebrafish neural crest cells. In the absence of evidence for the pigment cell-specific intermediates proposed before, we conclude that pigment cell fate specification proceeds directly from a broadly multipotent progenitor. We note that these HMP progenitors were identified at all stages examined, even at later stages (42-72 hpf) when NCC induction is thought to have long ceased. We hypothesise therefore that these HMP cells correspond to presumed satellite glia and Schwann cell progenitors readily detected using *sox10* expression in association with both ganglia and peripheral nerves^52–54^. We note that at least one population of these, associated with the DRG, has been identified as a population of adult pigment stem cells that generate the post-metamorphic pigment pattern of adult zebrafish^55,56^. Using an optimized RNAscope protocol we have demonstrated the unexpectedly early and widespread expression of *phox2bb* in premigratory and migrating NCCs, including in a well-studied population of putative Schwann cell precursors, providing the first evidence to our knowledge for a widespread autonomic neuron potential amongst zebrafish NCCs. It is likely our HMP progenitors correspond in mammals to the Schwann cell precursors that form the source of many adult melanocytes^29^; indeed, Schwann cell precursors have also been shown to be a source of parasympathetic neurons and adrenomedullary cells and hence have retained a broad multipotency ^57–60^.

We reconcile our new view of NCCS retaining high multipotency with the more conventional observations interpreted under a PFR model in former studies (e.g. ^13,14,23,61,62^) by noting technical differences in our approaches. Firstly, our use of NanoString technology and a refined RNAscope protocol has given increased sensitivity to low level gene expression. For example, expression of *ltk* in the progenitor cells in this study is rather variable, reflecting the notably low levels of WISH expression in the ‘early’ phases^13–15^, and consistent with the absence of *ltk* as a marker in recent scRNA-seq studies^61,62^. Our recent expanded WISH characterization identified co-expression with *ltk* of *sox10, tfec* and *mitfa* as characteristic of the putative chromatoblast (MIX)^13,14^, and we see these genes as more robust components of the ltHMP progenitor identified by our single cell transcriptional profiling here. Nevertheless, in our *ltk* fate mapping studies reported here the prolonged perdurance of GFP expression compensates for the low level and transient expression of *ltk* itself, allowing sensitive detection of the fate of *ltk*-expressing cells. Secondly, differences in single cell isolation protocols used may have acted alongside this elevated sensitivity. We note that our profiles for control pigment cells include unexpected expression of fate specification genes for other fates (e.g. *phox2bb*). Similarly in our studies of co-expression of *phox2bb* and *ltk*, although spatially widespread, only a subset showed overlapping expression *in vivo* even with our optimized RNAscope protocol. Together these observations suggest that the transcriptional profile of our single cells has ‘relaxed’, away from the apparently specified state seen in vivo by WISH, towards a more basal progenitor state during isolation. This provides a strong indication of the highly dynamic nature of the fate specification and differentiation process – that apparently specified cells identified *in vivo* are held in states biased towards one or more fates by environmental signals. It also provides evidence that these cells, even when strongly differentiated, were not yet committed at these stages. Thus, our protocol was apparently sensitized to detection of cell *potential*; in contrast, many former studies favoured detection of the state of fate *specification* of these cells.

To reconcile our new observations with those underpinning the PFR model, we propose a novel Cyclical Fate Restriction model in which our broadly multipotent intermediate is transcriptionally dynamic, transitioning between sub-states, each of which is likely biased towards a subset of derivative fates (Fig. 4d; ^42^). We propose that heterogeneity of HMP cells results from the intrinsically dynamic nature of their gene regulatory network (GRN). This would also explain the substantial heterogeneity of marker expression in premigratory neural crest cells that led to the PFR model (e.g. ^63–68^). These dynamic changes might be complex or irregular, but we favour the idea that they show a cyclical progression between states biased towards individual fates. This would reconcile key observations underpinning the PFR Model of zebrafish pigment cell development: why mutations that prevent the adoption of one specific fate (e.g. melanocyte (*mitfa*)) result in an increase in another specific fate (iridophore), since this would reflect the order in which GRN sub-states biased towards different fates are dynamically organized in the cycle^42^. It is becoming clear that a more dynamic view of stem cells may be required to understand their biology and we believe this will prove true for NCSCs too; sensitive direct or indirect readouts of GRN state transitions will be required to test this model in the future. We note that two of the progenitor clusters identified in our profiling (eHMP and ltHMP) show noticeably similar profiles, and speculate that these might represent two such sub-states; this idea is consistent with the main differences between the eHMP (low *ltk;* high *tfap2a, tfap2e, sox9b*) and ltHMP (high *ltk;* lower *tfap2a, sox9b*; low *tfap2e*) cell clusters identified here, but will require comprehensive experimental testing. These cells appear to persist into later embryonic stages; we speculate that these cells therefore include the persistent NCSCs deduced from studies of pigment cell regeneration and adult pigment cell development ^56,69,70^, but also that they include many other NCCs which *in vivo* occupy a variety of states of fate specification, with overlapping markers for 2 or more cell-types^13,14^, giving the impression of PFR. Interestingly, a detailed single cell transcriptomics study of mouse neural crest development showed broadly multipotent progenitor cells, with subsequent neural intermediates interpreted as bipotent progenitors^32^. However, the same study appeared to show markers of melanocyte potential (e.g. *Mitf*) and a Schwann cell marker (*Mbp*) widely distributed amongst many cells, suggesting a pattern consistent with the view we are proposing here. Taken together, our investigations of Ltk activity, NCC fate-mapping and single cell profiling show that zebrafish pigment cell progenitors show unexpected and robust multipotency, suggesting a new paradigm for neural crest development encompassing a novel dynamical view of multipotency that may resolve the long-standing controversies in this area. Such a view may be widely applicable in the context of other progenitor cell-types.

## Supporting information

Supplementary Table 3 TaqMan Primers

Supplementary Table 1 Nanostring Gene List & Expression SUmmary

Supplementary Tabel 2 MTE Primers

## Methods

### Ethics statement

This study was performed with the approval of the University of Bath ethics committee and in full accordance with the Animals (Scientific Procedures) Act 1986, under Home Office Project Licenses 30/2415, 30/2937 and P87C67227.

### Fish husbandry

Embryos were obtained from natural crosses and staged according to Kimmel et al. ^71^. All fish were housed according to FELASA recommendations^72^, in tanks filled with circulating system water at 28±0.2°C. System water is made up from Reverse Osmosis water, with synthetic sea salt added at an amount such that the conductivity of the water is kept at approximately 800uS. Average water quality data are as follows: pH: 7.30, general Hardness: 160mg/l CaCO3, Ammonia: 0 mg/l NH3, Nitrite: 0 mg/l NO2, Nitrate: < 10 mg/l NO3- and Conductivity: around 800µS/cm. Light cycle is 14 hours light/10 hours dark (Lights on at 08:00, off at 22:00 daily). All fish were in general good health and no specific diseases were observed throughout the colony. The fish were fed Paramecium from 5-15dpf, followed by Ziegler Larval AP100 powder from day 16 – 22dpf. From 23dpf onwards they are fed Sparos Zebrafeed and Brine Shrimp. Fish euthanized by schedule 1 killing were given an overdose of an anaesthetic (5 drops of 0.48% MS-222 added to a petridish of water).

### Construction of nucleophosmin-*ltk* fusion gene

Primers to amplify cDNAs corresponding to *NUCLEOPHOSMIN* (*NPM*) domain in human *p80* fusion gene ^48,73^ and to cytoplasmic kinase domain of zebrafish *ltk* gene ^15^ were as follows.

XbaI/EcoRI *NPM* upper : 5’-GCTCTAGAATTCATGGAAGATTCGATGGACATG

*NPM* lower SpeI : 5’-GACTAGTAAGTGCTGTCCACTAATATG

SpeI *ltk* upper : 5’-GACTAGTCTATTATCGTAAGAAGAACCACCTG

*ltk* lower XbaI : 5’-GCTCTAGAGCTACACAGGGGTGACACTCAG

*p80* cDNA for template of *NPM* was kindly provided by Dr Richard Jäger (Uniklinikum Bonn, Germany), and *ltk* cDNA is the one reported by our group ^15^. Amplified *NPM* and *ltk* fragments were ligated using SpeI site, and subcloned using XbaI site into modified pCS2+ vector, which contains *sox10-4725* enhancer ^74^ instead of cytomegalovirus (CMV) promoter, IRES sequence and membrane tethered form of *egfp* cDNA, to form *Tg(Sox10:NPM-ltk, egfp)*. As a negative control, we introduced a point mutation to exchange the conserved lysine with arginine ^48^ at 212 in ATP binding domain of the chimeric NPM-Ltk protein, to generate *Tg(Sox10:NPM-ltk_K943R, egfp)*.

### Combined whole mount fluorescent *in situ* hybridization and immunofluoresence

Anesthetised embryos were fixed in 4%PFA overnight at 4 °C. PFA was removed and 100% methanol was directly added. Samples were stored at −20 °C until processed. We used the RNAscope Multiplex Fluorescent kit V2 (Bio-techne, Cat No. 323110) following the manufacturer’s protocol with some modifications. Methanol was removed from samples and air-dried for 30 min at RT. 50 ml of Proteinase Plus was added for 10 min at RT and washed with 0.01% PBS-Tween for 5 min x3. Samples were incubated in 50 ml of hydrogen peroxide for 7 minutes at RT followed by 3 washes with RNAse free water for 5 min each. Samples were incubated overnight with diluted probes (1:100). Probes were recovered and samples were washed in 0.2X SCCT for 15 min x3. We followed the manufacturer instructions for AMP 1-3 and HRP C1-C4 using 2 drops of each solution, 100 μl of Opal 570 or 650 (1:3000) and 4 drops of HRP blocker. Washing in between these solutions was performed twice at RT for 10 min with 0.2X SSCT prewarmed at 40 °C. Samples were incubated in primary antibody rabbit a-GFP (Invitrogen Cat. No. A11122) diluted (1:750) in blocking solution (0.1% PBTween, Normal goat serum 5% and 1% DMSO,1:750:) overnight at 4 °C then washed 3x with 0.1% PBTween for 1 hour with agitation. Samples were incubated in secondary antibody Goat a-Rabbit Alexa Fluor488 (Invitrogen Cat. No. A32731TR) diluted (1:750) in blocking solution for 3 hours at RT and then washed 6x 30 min with 0.1% PBTween. Samples were counterstained with 2 drops of DAPI provided in kit for 3 minutes and then rinsed once with 0.1% PBTween. Samples were mounted in 50% glycerol/PBS in glass bottom petri dishes.

### DNA injection

Purified DNA construct was diluted to 50 ng/µl with water, and injected into cell body at 1-cell stage. Developing embryos were sorted and cultured at 28.5°C until the appropriate stage. To prepare embryos for in situ hybridization, we added phenylthiourea at 0.003%, whereas a quarter dose was used to allow them to be only weakly pigmented when we needed to detect fluorescent signals in melanocytes after antibody staining.

### Construction of *TgBAC(ltk*:*gfp)* and *TgPAC(sox10*:*gfp)* reporter

We identified a fully sequenced BAC clone containing long 5’ flanking region of *ltk* gene (CH211-254C11, 36100 bp, Accession number is CR387922.) in Zebrafish Information Network database, ZFIN (http://zfin.org/). The exon-intron structure of *ltk* gene in this BAC was determined by comparing our reported cDNA of *ltk* (accession number EU399812) with this BAC clone. This analysis showed that the BAC contained the coding sequence of the first exon in the area between 35072 bp and 35126 bp, though this coding sequence is longer than the one in the first exon reported in our previous paper^15^. To insert the *gfp* cDNA into this BAC, we used BAC recombineering^49^. The targeting vector, containing *egfp* cDNA with bovine growth hormone polyA and *kanamycin resistant* gene as a selection marker^50^, and bacteria (SW101) used for recombination were obtained from Dr. Higashijima and from Biological Resources Branch of Frederick National Library for Cancer Research, respectively. We prepared fragment to be inserted into BAC by high fidelity PCR using targeting vector as template. The primers were as follows.

*ltk*-5arm :

5’-AGAGATTAGGCTAACAAACACTTTATCTCCGGGATCCTTTTTAAGGAGCC-atggtgagcaagggcgagga

*ltk*-3arm :

5’-TTAACGTTAACAGAAACCAGCAGGCCAGTATTAATTAGCAAAACACTCAC-cagttggtgattttgaactt

The sequences in lower case of *ltk*-5arm and *ltk*-3arm target the *gfp* cDNA and *kanamycin resistant gene*, respectively. The sequences in upper case correspond to the *ltk* genomic sequence, and act as homologous domains during recombination. The recombineered resultant was confirmed by PCR targeting the two BAC ends and two junctions between *ltk* gene and *egfp* or *kanamycin resistance* gene. To make transient transgenic embryos, we simply injected 100 pg complete construct into fertilized eggs.

As a positive control construct to demonstrate how readily each NCC derivative was labelled when GFP was expressed in early NCCs, we recombined gfp into an 84 kb P1 phage-derived artificial chromosome (PAC) containing the zebrafish sox10 gene and 30 kb upstream and 52 kb downstream sequences (clone BUSMP706I16137Q2). To create this TgPAC(sox10:gfp) construct, we used RecA-dependant recombineering ^75^ as described in ^76^. Characterization of TgPAC(sox10:gfp) transient transgenics confirmed that it accurately reproduced the early sox10 expression pattern, labelling all premigratory NCCs ^76^, and thus behaved similarly to a plasmid construct containing 4.9 kb of upstream sequence characterized previously ^52,74^.

### Analysis of cell fate in transient transgenic fish

The derivatives of NCCs examined were pigment cells (melanocytes, xanthophores and iridophores), glial cells (satellite glia in DRG, Schwann cells around the posterior lateral line nerve), neurons (sensory in DRG, enteric and sympathetic neurons) and jaw cartilages. All were examined sequentially in injected embryos as shown in Fig. 4B, using bright field, Nomarski or fluorescence optics as appropriate on a Nikon Eclipse E800 or Zeiss Confocal 510 META. Rohon-Beard neurons and posterior lateral line ganglia were also labeled in injected embryos. These, like other neurons, were counted after immunostaining with anti-GFP/Hu antibodies. The morphological criteria used for these cell types were as follows. 1) Melanocytes: Black pigment, dendritic form and location on neural crest migration pathways or in stereotyped locations. 2) Xanthophores: Yellowish color, granular pterinosomes, dendritic shape, and peripheral location immediately under epidermis. 3) Iridophores: localisation in dorsal, ventral (including lateral patches) or yolk sac stripes, and their round shape. 4) Satellite glia: GFP positive cells without Hu signal in DRG. DRG position was confirmed by *neurogenin1*(*ngn1*) promoter-driven red fluorescent protein (RFP) using *Tg(−8.1ngn1:RFP)*^51^. 5) Schwann cells: Elongated shape and GFP signal around the posterior lateral line nerves and spinal nerves; position of nerves was confirmed by anti-acetylated tubulin antibody. Identification further confirmed by noting that GFP signal and anti-acetylated tubulin signal seemed interwoven with each other. Cell-types 4) and 5) given as glia (gli). 6) DRG neurons: *ngn1*-driven RFP in typical position next to the spinal cord. 7) Enteric neurons: Hu signal localised to gut surface. 8) Sympathetic neurons: Hu signal localised to area between ventral stripe and notochord. 9) Lower jaw cartilage: cuboidal shape and position in jaw. We counted only the cells that have RFP or Hu as neurons. Some other cells appeared likely to also be neural crest-derived neurons, such as enteric or sympathetic neurons, because of their position, but these were not included here unless marker was positive. However, neuron marker negative cells in DRG position were counted as “DRG” because they were considered to be either undifferentiated neural precursors or satellite glia, which are both neural crest-derived, but which could not be distinguished in our study. These transient transgenic studies of *in vivo* potency of *sox10* and *ltk-*expressing cells were performed as follows. We first scored live embryos at 2 dpf for the presence of GFP fluorescence in DRG neurons by assessing overlap with RFP signal; at 3 dpf we scored for GFP expression in melanocytes, identified by their endogenous pigment; finally at 4 dpf, we scored xanthophores by the granular appearance of their pigment granules as observed with DIC optics, and fixed the embryos and processed them for morphological observation and for immunofluorescence using anti-GFP, anti-Hu, and anti-acetylated tubulin antibodies to confirm the presence of other NC-derived cell-types expressing GFP (iridophores, enteric neurons, sympathetic neurons, Schwann cells and cartilage).

### FACS collection of single neural crest cells

We used embryos with following genotypes for single cell profiling: (1) wild type AB Zebrafish line, (2) *Tg(Sox10:Cre)^ba74^xTg(hsp70l:loxP-dsRed-loxP –Lyn-Egfp^tud107Tg^* transgenic fish line, and (3) *Tg(Sox10:Cre)^ba^*^74^*;Tg(hsp70l:loxP-dsRed-loxP –Lyn-Egfp^tud107Tg^;sox10^m618/m618^*.

### Cell Dissociation

Embryos from required stock were grown up to the desired stage (from 14 to 72 hpf) in standard embryo media at 29° C. To prevent melanisation in embryo melanocytes, PTU *(N*-Phenylthiourea, Cat# 7629, Sigma-Aldrich) was added at a final concentration of 0.003% at 24 hpf. To stimulate eGFP expression, embryos were heat-shocked by placing them in 42°C embryo media followed by 1 hour incubation at 37° and at least 1 hour incubation at 29°C. If required, the embryos were dechorionated using pronase (Pronase from *Streptomyces griseus,* Cat# 000000010165921001, Sigma-Aldrich) at a final concentration of 1 mg/ml^77^. The heads were cut from all the embryos at stage 30 hpf or older to decrease the number of *sox10*-positive cells of craniofacial skeletal and otic fates. Embryos were then digested as previously described with small modifications ^78^. In brief, embryos were rinsed with Ca-, Mg-DPBS (Sigma, D8537), placed in a flask containing TrypLE™ Express Enzyme (Cat#12605036, ThermoFisher Scientific) in ratio of 10 ml per 100 embryos, containing 0.003% Tricaine; incubated for 30-90 min at 100 rpm, 37° C in the shaker incubator with constant monitoring until the embryos were digested to a mixture of single cells and small fragments of tissue; then digestion mixture was triturated 10-15 times, using a Pasteur pipette; passed through 100-micron strainer (MACS SmartStrainers, Cat# 130-098-463, Miltenyi Biotech.) into 50 ml Falcon tube and centrifuged for 5 min, 500xg, 4° C. The cell pellet was re-suspended in DPBS and the cell suspension was passed through 30-micron strainer (MACS SmartStrainers, Cat# 130-110-915, Miltenyi Biotech.) into 50 ml Falcon tube and centrifuged again for 5 min, 500xg, 4° C following by re-suspending the cells in 0.5-1 of ml cell isolation media (2% FCS, DPBS:HBSS=1:1 and 1mM SYTOX Blue Dead Cell stain (ThermoFisher Scientific). Cells were imaged before FACS and after FACS to confirm successful purification of GFP+ cells.

### Single cell sorting of eGFP-positive cells into 96-well plate

Single eGFP-positive cells were sorted into each well of 96-well plate containing the lysis buffer. For cell lysis and further cDNA synthesis and Pre-Amplification we used the protocol supplied by Bio-RAD (http://www.bio-rad.com/webroot/web/pdf/lsr/literature/Bulletin_6777.pdf) with slight modifications according our primer design, and with very reproducible results. The lysis buffer, was prepared on ice according the protocol, using the whole SingleShot™ Cell Lysis Kit (SingleShot™ Cell Lysis Kit, 100 x 50 µl reactions Cat# 1725080, Bio-Rad) as follows: 0.8 ml SingleShot Cell Lysis Buffer, 0.1 ml Proteinase K, 0.1 DNaseI ml, 4 ml TE buffer (Sigma-Aldrich). Spike-in RNA (polyadenylated kanamycin mRNA #C1381, Promega, USA) was added to the Cell Lysis Buffer with a final concentration 10^7^ molecules/ml. 4 μl of cell lysis buffer was aliquoted into 96-well semi-skirted PCR plates on ice and immediately frozen at −80°C. AriaIII Cell sorter was set with optimal flow parameters, the whole system cooled to 4o C, drop delay appropriately adjusted and the precise positioning of the home device for the 96-well plate. Cells were excited with 488 nm laser and gated using forward and side scatter to avoid debris and smallest particles. SYTOX™ Blue Dead Cell stain (ThermoFisher Scientific) was added to final concentration of 1uM. Viable SYTOX™ Blue-negative; GFP-positive cells (P2) were gated using the 405 nm laser and 450/50 nm filter (DAPI channel) and 488 nm laser and 450/50 nm filter (GFP/FICT channel). Cells were sorted using 488 nm laser and 450/50 nm filter (GFP/FITC channel), and the 561 nm laser and 582/15 nm filter (DsRed channel with thresholds established by comparison with stage-matched non-transgenic AB wild type fish, selecting the discrete population of bright GFP-positive cells and excluding bright red cells to avoid auto-fluorescent particles (see Supplementary Methods Supplementary Figure 1). Cells were sorted into each well of the plate using “single cell” setup, the plate was immediately placed on ice, sealed, vortexed for 10 seconds, centrifuged briefly and underwent genomic DNA digestion by placing the plate in a thermocycler and incubating at 25°C for 10 min and 75°C for 5 min, followed by holding at 4°C.

### Single cell sample preparation and nCounter (NanoString Technologies) quantitation of gene expression

Total RNA was converted to cDNA, using iScript™ Advanced cDNA Synthesis Kit for RT-qPCR (Cat# 1725038, Bio-Rad) following by 25 cycles of preamplification using mix of 47 pairs for MTE primers, each at final 50nM concentration (Supplementary information, Supplementary Table 2) and SsoAdvanced™ PreAmp Supermix (Cat# 1725160, Bio-Rad). After amplification, samples were checked for quality control, and those which passed QC were selected into 12-samples strips and stored at −80°C. When required, pre-amplified samples were thawed, hybridized with both Reporter and Capture probes, applied to the chip and chips were analysed using nCounter according to the manufacturer protocols (NanoString Technologies) https://www.nanostring.com/applicatio/files/4714/9264/4611/nCounter_XT_Assay_Manual.pdf. The NanoString data were recorded as DNA molecule counts for each gene of interest, and then were subjected to further statistical analysis.

### Quality control of the pre-amplified samples

To check absence/presence of a cell in each well of 96-well plate of sorted cells, and to check the efficiency of the preamplification steps, we utilised two custom designed Quality Control Taqman Assays: housekeeping gene *rpl13* Assay and in-RNA spike control Kanamycin Assay. Primers and probes for both assays were designed to recognise the amplicons used for preamplification with MTE primers (Primer3 Plus software, (http://www.bioinformatics.nl/cgi-bin/primer3plus/primer3plus.cgi).

The following primers and probes were used:

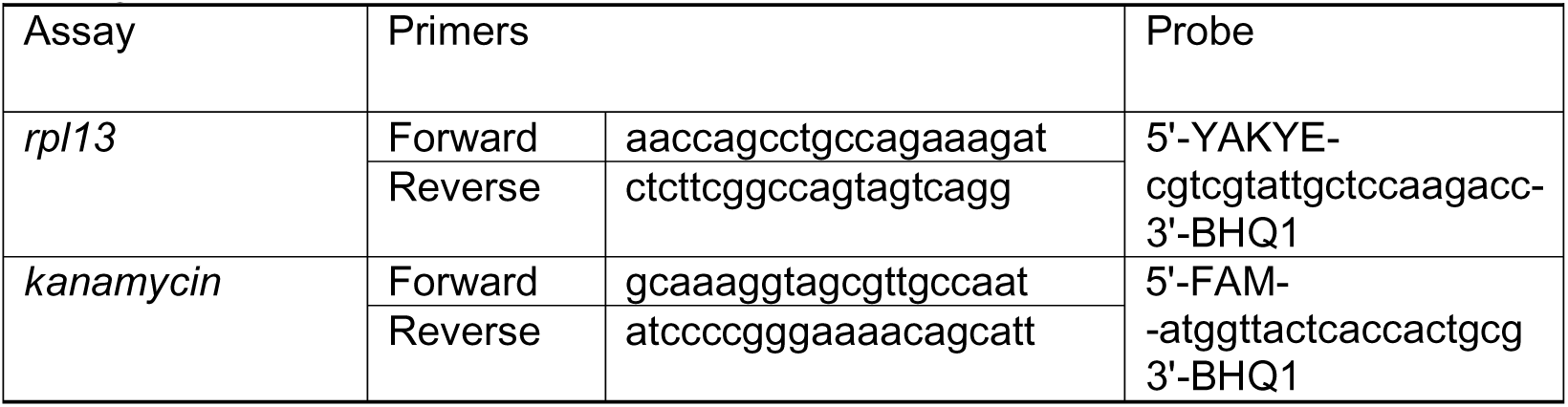

Real time quantitative PCR was performed with TaqMan™ Fast Advanced Master Mix, (#4444965, ThermoFisher Scintific) according the manufacturer’s protocol using StepOnePlus™ Real-Time PCR Systems (Applied Biosystems). Ct values were estimated using an automatic baseline and a standard threshold of 0.2. For *kanamycin*, the samples demonstrating Ct-values less than 14 were rejected as poorly amplified, for *rpl13*, the samples with Ct-values less than 24 were rejected as those with poor RNA quality or empty.

### Pigment Cell Enrichment

72 hpf embryos were processed as previously described^78^ with minor modifications. Briefly, fish were anesthetized with Tricaine, decapitated and digested using TrypLE Express 10 ml per 100 embryos. After incubation for 45-120 min at 37° C while shaking at 100 rpm, the suspension was triturated with a Pasteur pipette. Dissociated cells were filtered through a 300 micron strainer and immediately through a 70 micron strainer to the 50 mL Falcon tube, precipitated at 500 g for 5 minutes at 4°C, resuspended in 1 mL cold isotonic Percoll (Sigma, P1644), transferred to 1.6 mL Eppendorf tubes and spun at 2000 rcf for 5 minutes at 4°C in an angle rotor (Thermo Scientific Fisher Heraeus Biofuge Primo R Centrifuge). The pellet containing pigment cells was resuspended in 400 µL of ice cold Cell Media (CM, DPBS:HBSS(no calcium, no magnesium, Cat #14170112, ThermoFisher Scientific) = 1:1, supplied with 2% fetal Bovine serum (heat inactivated, Cat#10082139, ThermoFisher Scientific), and placed onto preformed Percoll density gradients prepared as previously described^78^. The cells were centrifuged in a swinging bucket rotor at 2000 rcf for 10 minutes at 4°C following by re-suspending in Cell Media and centrifugation at 500g for 5 min. Pellet was re-suspended again in 0.5-1 ml of Cell Media, and kept on ice until FACS.

### Selection of melanocytes and iridophores from pigment cell enriched suspension

We utilised the natural physical properties of melanocytes and iridophores to absorb and reflect light respectively and sorted them from the suspension of pigment cells using Fluorescence-Activated Cell Sorting (FACS). Pigment cells enriched after Percol gradient centrifugation were resuspended in CM and filtered through 30-micron strainer (MACS SmartStrainers (30 µm), Cat# 130-110-915, Miltenyi Biotech.). Cells were sorted with an AriaIII cell sorter using a 130-micron nozzle, in sterile conditions at +5°C, with optimized flow parameters, drop delay and correct positioning of the 96-well plate. Cells were excited with 488 nm laser and gated using forward and side scatter to avoid debris and smallest particles. Cells were sorted based on autofluorescence using 488 nm and 561 nm filters, corresponding to DsRed and GFP channels (Supplementary Methods Supplementary Figure 2). Iridophores were selected by high excitation in both channels, forming a scattered population in the upper right quadrant. Melanocytes do not possess any fluorescence and were detected at the lower left of the quadrant of the DsRed-FITC plot. Cells were collected into ice-cold LB buffer in 96-well plates and immediately processed for genomic DNA degradation followed by cDNA conversion, and kept at −80°C until next step as described above. Alternatively, cells were directly sorted to a tube containing 0.5 ml of Trizol reagent (TRIzol™ Reagent, Cat:# 15596026, ThermoFisher Sci.) and kept on ice until mRNA extraction.

### mRNA Extraction and cDNA conversion

To extract RNA from Trizol stocks we used Direct-zol™ RNA MicroPrep (Cat.# R2062, ZymoResearch, USA) according the manufacturer’s protocol. The total RNA was extracted in 10 ml, measured using Nanodrop-2000, and tested with Experion™ RNA HighSens Analysis Kits (Cat.# 700-7105, Bio-Rad) using the Experion™ Automated Electrophoresis System (Bio-Rad). 5 ml of total RNA from either melanocytes or iridophore (50-60 ng/ml) was used directly for NanoString expression analysis.

cDNA was synthesized using 1 μl of total RNA from melanocytes or iridophores with iScript™ Advanced cDNA Synthesis Kit (#1725038, Bio-Rad) following by 6 cycles of pre-amplification with pooled 47 pairs of MTE primers designed for NanoString CodeSet (SsoAdvanced™ PreAmp Supermix, Cat# 1725160, Bio-Rad); amplified dsDNA was analysed using the NanoString expression protocol.

### Single Cell Quantitative RT-PCR profiling

Primers and probes for TaqMan qPCR expression assays were designed using the Primer3 web-site and ordered from Eurofins (http://www.eurofins.com/genomic-services/). Amplification primers and TaqMan assays are represented in Supplementary information (Supplementary Table 3).

Zebrafish transgenic embryos at stage of 30 hpf were decapitated; eGFP-positive cells were FAC-sorted and RNA was amplified as described above with the set of 15 pairs of amplification primers (Supplementary Table 3). Reaction mix was diluted twice with nuclease-free water (Nuclease-Free water, Cat# P119C, Promega) and 1 ul of amplified dsDNA was used for gene specific TaqMan Assay using the manufacture protocol for TaqMan™ Fast Advanced Master Mix (Cat# 4444964, ThermoFisher Scientific). Expression data was normalized relative to *kanamycin* for each sample, filtered to eliminate samples with low gene expression, and transferred to R Seurat package for further analysis (see Supplementary methods). Heatmaps were built with ComplexHeatmap R package. For original data, see Supplementary Table 4.

#### Single cell data analysis

We followed general recommendations of Luecken & Theis^79^ for our single cell data analysis protocol. For the detailed description of the data and data processing protocol see Supplementary Methods text.

### The initial data

The results of NanoString transcriptome contained 1090 cells of regular and control cell types, including WT cells and sox10-/- mutants. Supplementary Methods Supplementary Table 1 contains cell type statistics for the initial set.

#### Quality filtering of the cells

To ensure the high quality of gene expression measurements we used a series of filters, test the data set for reliance of the control probes and internal consistency. We removed cells with poor total counts of test probes other than those of housekeeping genes (*rpl13* and *kanamycin*), assuming that most of them related to cell types expressing other cell markers than those represented in our panel. In addition, we removed data for probes (genes) that displayed expression in only a small number of cells. We used the NanoStringNorm R package^80^, and removed a number of cells with poor norm factors and noisy background. Normalization was performed using the sum of probes (kanamycin spike-in and *rpl13* internal controls) as the reference housekeeping class with ‘mean and 2sd’ selected for the background. In total 731 cells (25 control iridophores, 19 control melanocytes, 108 cells from tails, 444 regular WT cells from different stages and 135 *sox10* mutant cells) survived filtering and normalization. We filtered the expression matrix nullifying elements with values less than 30 and imputed for dropouts using drImpute^81^. In total about 20% of zero counts have been imputed into meaningful quantitative values. After imputation the log transformed expression values have been loaded into a Seurat object (ver. 2.3.4)^82^.

#### Descriptive statistics

Principal component analysis demonstrated that control melanocytes, control iridophores and tail cells were shifted from the center of the main cloud in some projections (Supplementary Figure 4). For example, melanocytes were clearly shifted along the PC2 component, with high weights of melanocyte gene markers (*mlphb, slc24a5, oca2, tyrp1b, sylva*)^78^ forming a clearly separated cloud, which included also some regular cells, apparently NCC derivatives differentiating into melanocytes. In contrast *sox10* mutant cells (Supplementary Figure 4, coloured in magenta) were not offset from the central cloud of WT cells, which is consistent with the previous suggestion that mutant cells are ‘trapped’ in a progenitor state (Dutton et al 2001). We observed that regular cells from different stages (hpf) did not display significant differences in their distribution, and decided to consider them as a single ‘regular’ class.

#### Dimension reduction

We used UMAP and tSNE, two independent algorithms for non-linear dimension reduction which conserve intercell distances in high dimensions (42D space of gene expressions). Cosine distance implemented in R package `proxy` was used. The control cell types were even better separated in 2D UMAP and tSNE planes (Supplementary Figure 8); *sox10* mutants were not separated from the main cloud of WT cells overlapping with location of tail control cells but not overlapping with melanocyte and iridophore clouds (Extended Data Figure 1b).

#### Cell Clustering

We clustered cells using as features their gene expression profiles after dimension reduction used sharing nearest neighbor clustering algorithm (Waltman et al., 2013) implemented in the Seurat toolbox. Since UMAP is a non-linear transform which can sometimes bring about irrelevant cluster structure we used an enumerative algorithm to test all combinations of UMAP transform parameters and cluster resolution (each combination of parameters was tested three times). To evaluate the clustering quality we used control melanocytes and iridophores optimizing the proportion of the control cells found in the single cluster. Clusters with similar gene expression profiles were merged using `Seurat:ValidateClusters` procedure based on 4 top marker genes, arriving at 7 distinct clusters (Figure 2c). Given the stochastic nature of UMAP transformation it was encouraging to obtain robust clusters with same values of transformation/resolution parameters reproduced in replicates. Clusters containing control cell types (melanocytes and iridophores) and xanthophores (X) (identified by high expression of *pax7a*, *pax7b* marker genes) never merged with other clusters up to very high `ValidateCluster` threshold.

#### Pseudotime ordering

We used `slingshot` software^40^ to construct pseudotime trajectories in the 42D gene expression space. We required trajectories to begin from eHMP and end at melanocytes and iridophores (Supplementary Figure 8). The paucity of markers of xanthophores explains why we were unable to obtain a relevant trajectory to xanthophores.

#### Heatmaps

We used ComplexHeatmap package^83^ to construct heatmaps. Cosine distance implemented in R package `proxy` was used to cluster gene expression rows and columns when required.

#### Data availability

The Nanostring nCounter raw data and TaqMan assay data that support the findings of this study are available in Zenodo with the identifier DOI: 10.5281/zenodo.4953911 in the folder SourceData. The data are distributed under CC-BY licence.

## Code Availability Statement

Computer code is available in Zenodo with the identifier DOI: 10.5281/zenodo.4953911 and distributed under MIT licence. The code has been verified for its compliance with guidelines in Nature Code and Software submission checklist.

## Acknowledgements

The authors gratefully acknowledge the Technical staff within the Department of Biology & Biochemistry at the University of Bath for technical support and assistance. We gratefully acknowledge Nathaniel S. Gray for providing us with ALK inhibitor, TAE684, Dr Richard Jäger for *p80* cDNA for template of *NPM*, reagents for BAC recombineering from Biological Resources Branch of Frederick National Library for Cancer Research, and Dr Shin-ichi Higashijima for *egfp*-*polyA* cassette for recombineering. Dr Rosalind John (University of Cardiff) kindly supplied the reagents for the RecA mediated PAC recombineering. We thank Prof. Alfonso Martinez-Arias for helpful discussions in the early years of this project. We thank our colleagues Profs Adele Murrell and Andrew Ward for their critical comments on an earlier draft of this manuscript. This work was supported by Uehara Memorial Foundation (MN), Wellcome Trust VIP awards (MN), and BBSRC grants BB/ L00769X/1(RNK,HS,TS,HS) and BB/S015906/1 (RNK,JHPD,KCS,GB) and BB/L007789/1 and BB/S01604X/1 (AR), National Natural Science Foundation of China, Grant Number: 31000542 (XY), Royal Society International Exchange Cost Share 2017 Russia award (RNK) and Russian Fund of Basic Researcher grant 17-54-10014\19 (VM), University of Bath PhD Studentship and ORS award (TJC).

## Author contributions

RNK, HS, JHPD and AR conceived and designed the study and obtained funding. MN, TS, KCS, GB, XY, FSLMR and TJC performed the experimental studies and analysed and interpreted the data obtained. ASK and VJM designed the pipeline for Nanostring data processing, LAU and VJM wrote the code. RNK and VJM drafted the manuscript, and all authors contributed to revision of the manuscript. All authors have approved the submitted version.

## ADDITIONAL INFORMATION

**Supplementary information** is available for this paper.

Reprints and permissions information is available at www.nature.com/reprints.

## Extended data figure/table legends

**Extended Data Fig. 1.**
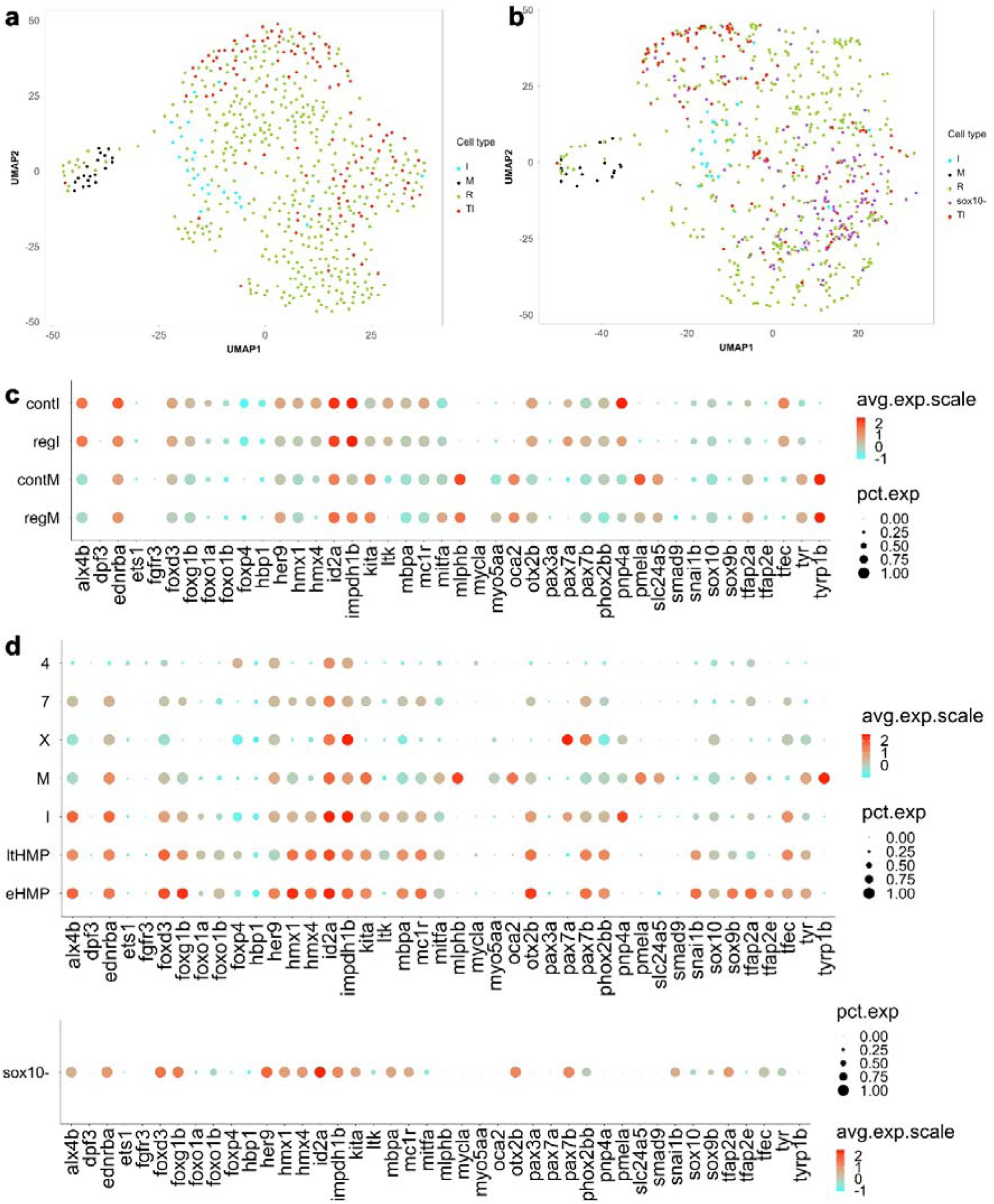
Distribution of control cells across clusters. **a**, Control melanocytes (M), control iridophores (I) and control tail cells (24 hpf)(Tl), plotted on UMAP projection of all regular WT cells (R). **b**, 2D UMAP projection shows that *sox10* mutants (sox10-; purple) occupy a region overlapping with some of the WT HMP cells (clusters enriched with Tl cells). **c**, Dot plot comparison of gene expression profiles of control pigment cells and regular cells found in the same clusters. Melanocytes: control cells (contM) and others from FACS-sorted WT population (regM). Iridophores: control cells (contI) and others from FACS-sorted WT population (expI). Highly expressed genes in each cell-type match expectations from published literature. **d** Dot plot of gene expression profile of WT clusters (see Fig. 2c) and *sox10* mutant cells. Expression colour scheme reflects expression of all genes and is not gene specific.

**Extended Data Fig. 2.**
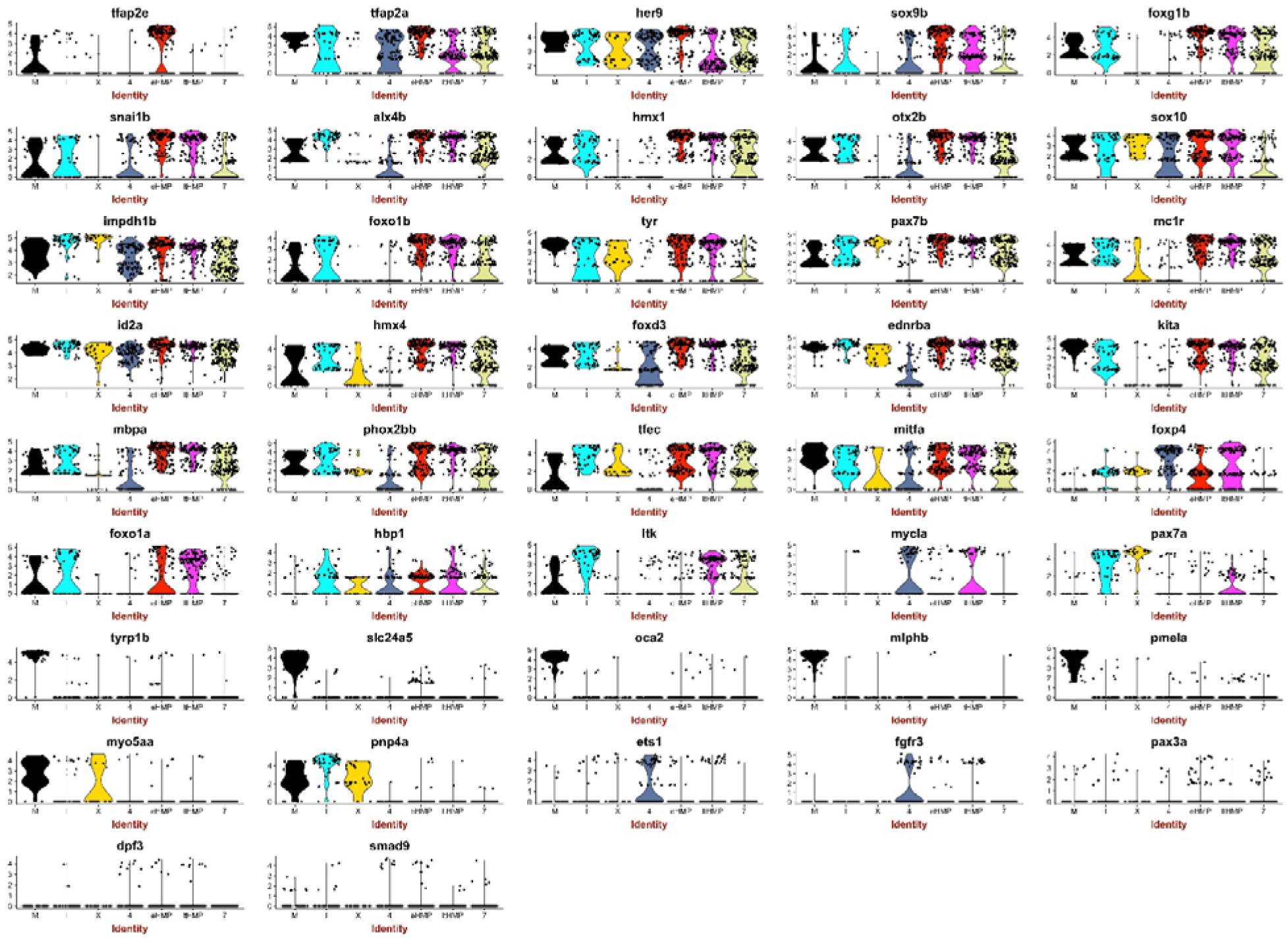
Gene expression profiles of WT cell clusters (see Fig. 2c). Violin plots of expression levels of each gene in each WT cell.

**Extended Data Fig. 3.**
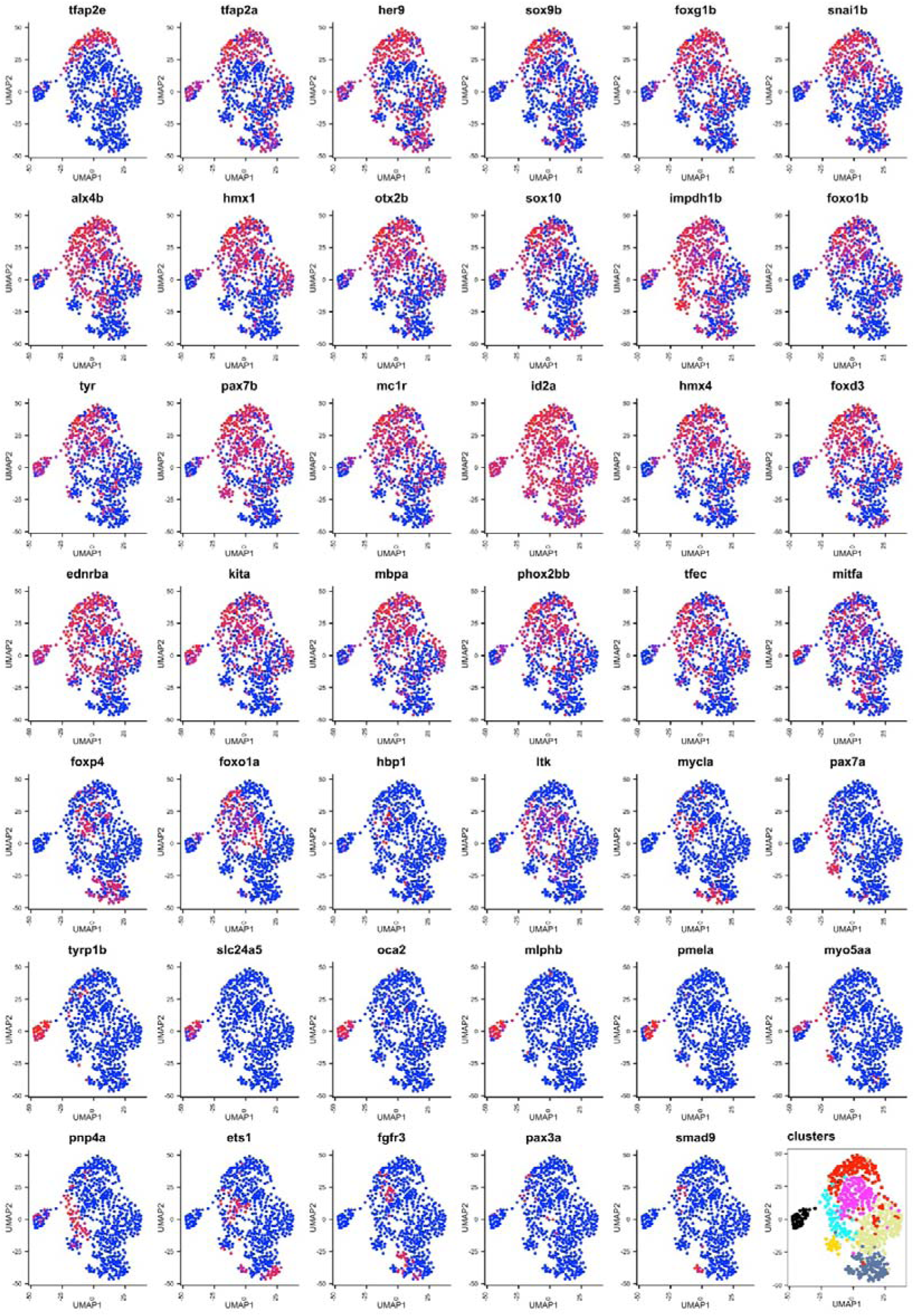
Gene expression profiles of WT cell clusters. Feature plots show regionally localized gene expression in different areas of 2D UMAP plane. Early neural crest specification markers (*sox9b* and *snail2,* but also *tfap2a* and *tfap2e*) are expressed in the cells found in the top right corner of the plane (eHMP and ltHMP clusters). Other genes like *ltk* and *foxo1a* are expressed more centrally (ltHMP). Note how eHMP and ltHMP clusters express genes with known role in fate specification of all pigment cell (*mitfa, pax7b, tfec, ltk)*, neuronal (*phox2bb)* and glial (*sox10*) cell-types, whereas no clusters show specific combinations consistent with MIXG (*mitfa, pax7b, tfec, ltk, sox10,* but not *phox2bb*) nor MI (*mitfa, tfec, ltk,* but not *pax7b* or *phox2bb*) intermediates. Specific markers of differentiating melanocytes (including *pmel, mlphb* and *oca2*) readily identify the discrete melanocyte cluster on the left. The iridophore cluster is less distinctive due to the known early progenitor expression of many key iridophore specification genes, but lies to the left of the centre in these plots and shows concentrated expression of *pnp4a, ltk, tfec, ednrba.* This analysis also confirms our recent demonstration from whole mount in situ hybridization studies that *pnp4a* is widespread in pigment cells (Petratou 2018). Bottom right panel shows clusters.

**Extended Data Fig. 4.**
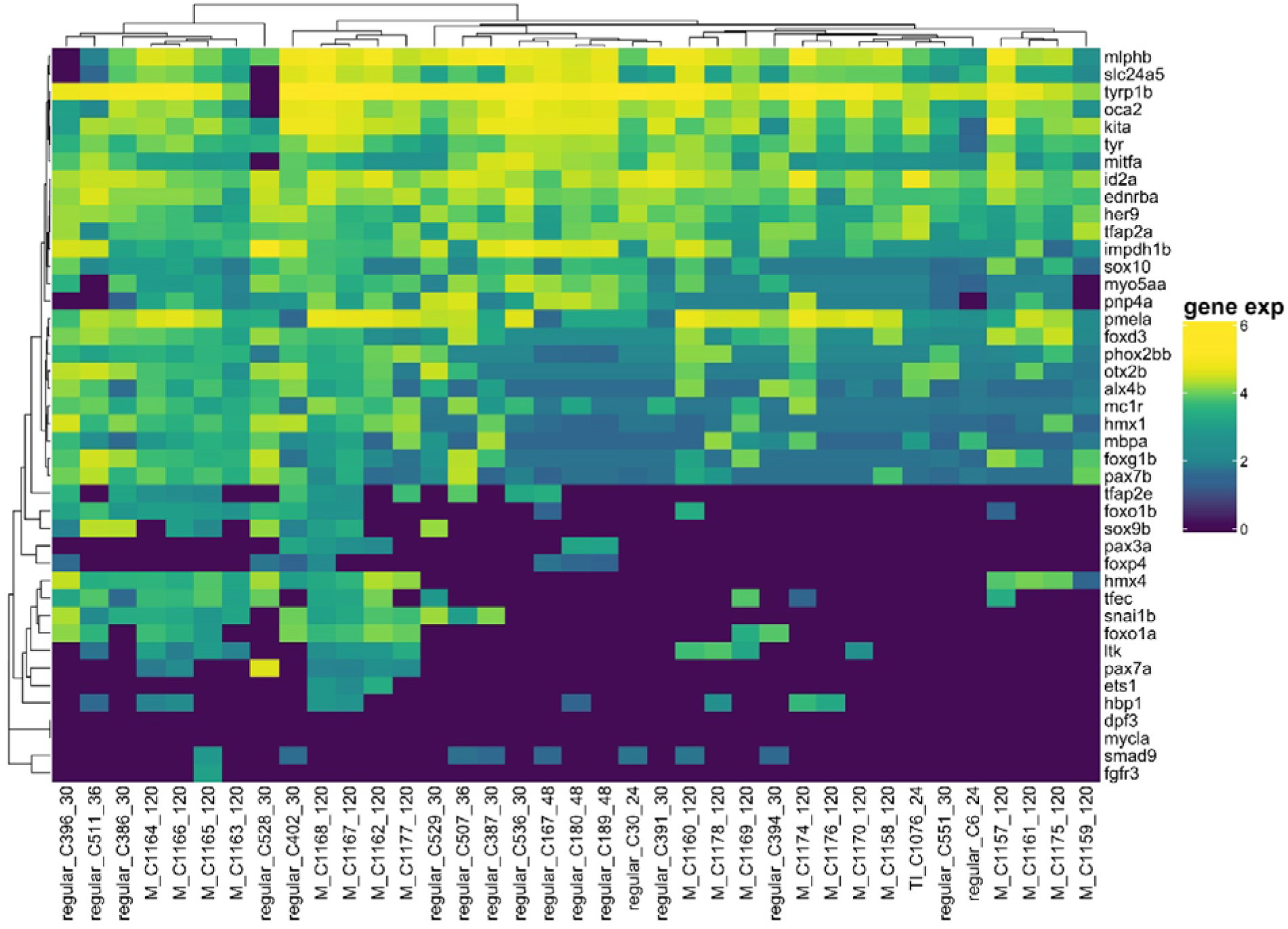
Melanocyte cluster contains two distinct sub-clusters. Expression profiles of melanocytes shows distinct sub-clusters; cells in right clusters show detectable expression of genes (e.g. *tfec, ltk, sox9b, snail2*) associated with earlier stages. Cell labels distinguish control melanocytes (labels begin ‘M_’) from those identified from regular WT cells (labels begin ‘regular_’) and from control Tail cells (labels begin ‘Tl_’).

**Extended Data Fig. 5.**
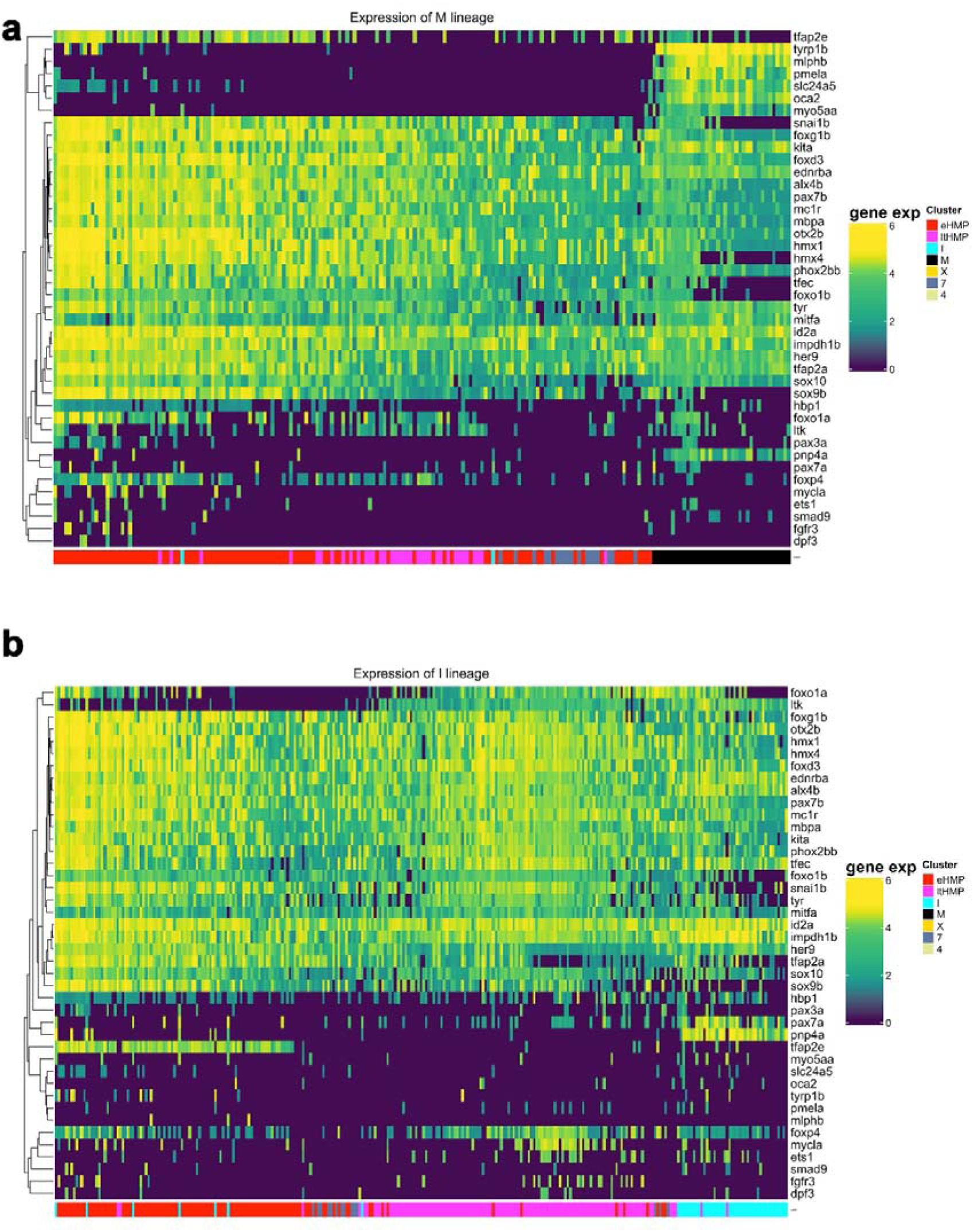
Pseudotemporal analysis of pigment cell differentiation. Heat maps showing pseudotime differentiation trajectories for melanocytes (**a**) and iridophores (**b**). Note that these representations were generated by collecting cells close to the trajectories drawn through the centres of clusters ordered into the cluster lineage tree in 42D space of gene expression. The trajectories (principle curves) were also constructed in 42D gene expression space and used to order cells in the heatmap. The cells in the colour chart below have been coloured according to assignment to particular clusters constructed after dimension reduction with UMAP to 2D.

**Extended Data Fig. 6.**
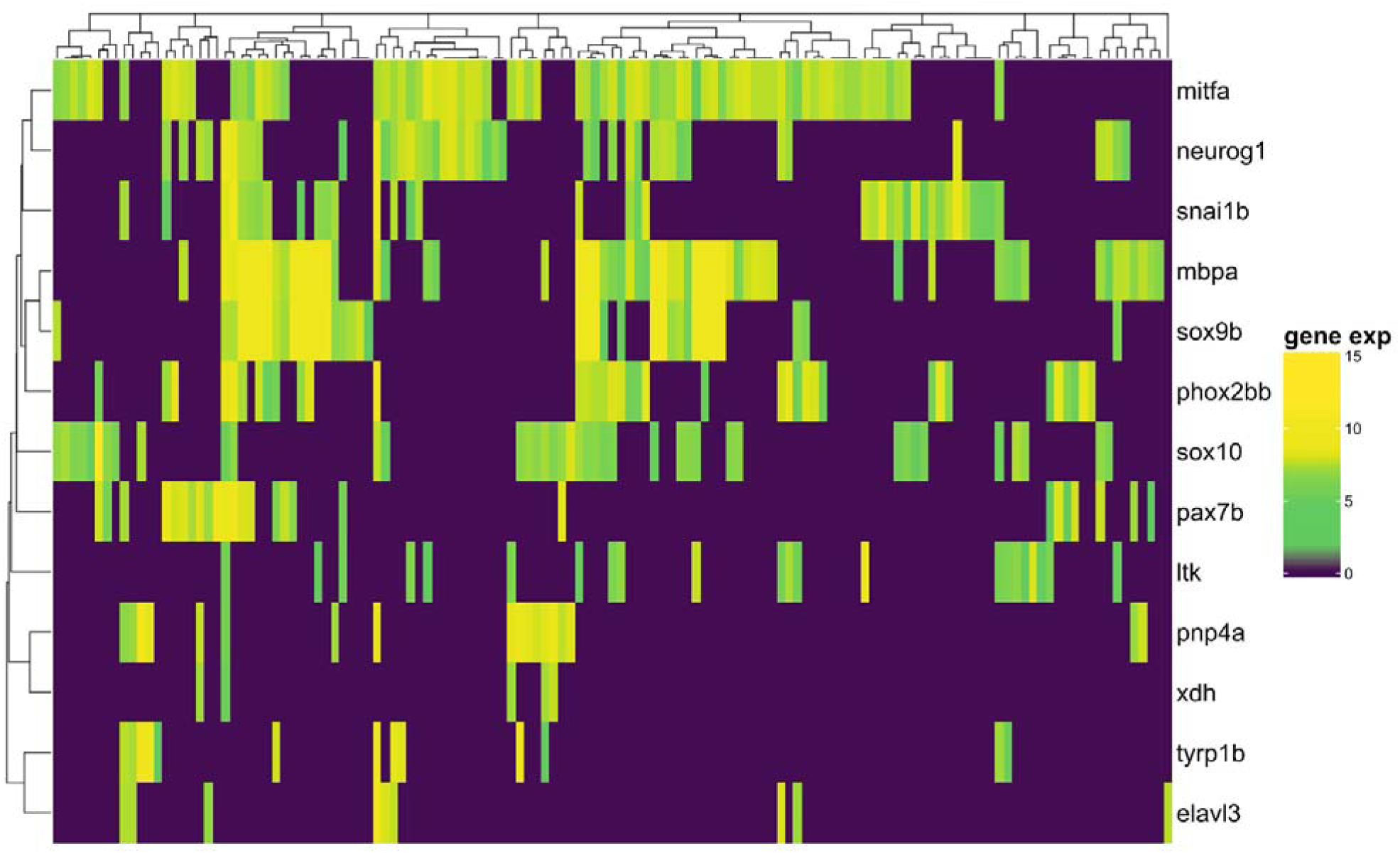
Independent TaqMan assay qRT-PCR assessment of overlapping fate specification gene expression in 24 hpf WT embryos. Note that individual cell profiles often show expression of fate specification transcription factor genes (*mitfa, pax7b, tfec, ltk, sox10, phox2bb*), with multiple genes being detected in each individual cell, whilst usually lacking detectable expression of pigment cell and neuronal differentiation markers (*pnp4a, xdh, tyrp1b, elavl3)*. Note also widespread expression of glial differentiation marker *mbpa*.

**Extended Data Fig. 7.**
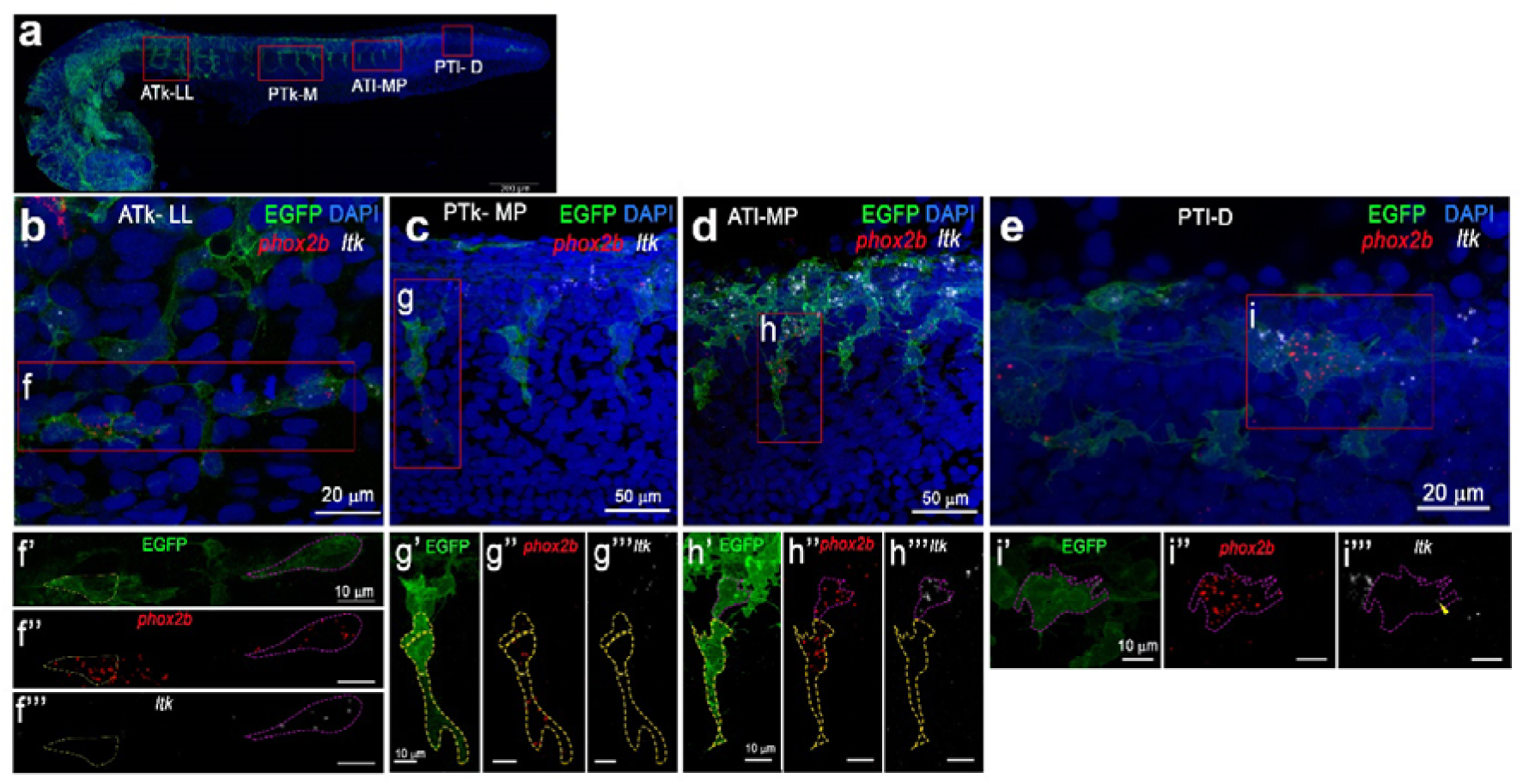
Co-expression of *phox2bb* and *ltk in vivo* by RNAscope. Simultaneous immunofluorescent detection of EGFP and RNAscope FISH for *phox2bb* (red) and *ltk* (white) on 24 hpf *Tg*(*sox10:cre; hsp70I:loxp-dsred-STOP-loxp-LYN-EGFP*) embryos after brief heat-shock. **a**, Projection of lateral views shows the lateral line (LL) in the anterior trunk (ATk; **a**), medial pathway (MP) in the posterior trunk (PTk; **b**), medial pathway (MP) in the anterior tail (ATl; **c**) and Dorsolateral (D) Posterior tail (PTl; **d**); NB this is same image as shown on panel **a** in Fig. 2. Red boxes indicate close-ups (**b-e**), for which single channel images of EGFP, *phox2bb* and *ltk* for insets boxed in **b-e** are shown in **f’-i’’’**. Cell membrane border of EGFP positive cells double-labelled with *phox2bb* and *ltk* are indicated with magenta dashed lines, whereas cell membrane border of *phox2bb* positive cells (lacking *ltk*) are indicated with yellow dashed lines. Scale bar dimensions are as indicated.

**Extended Data Fig. 8.**
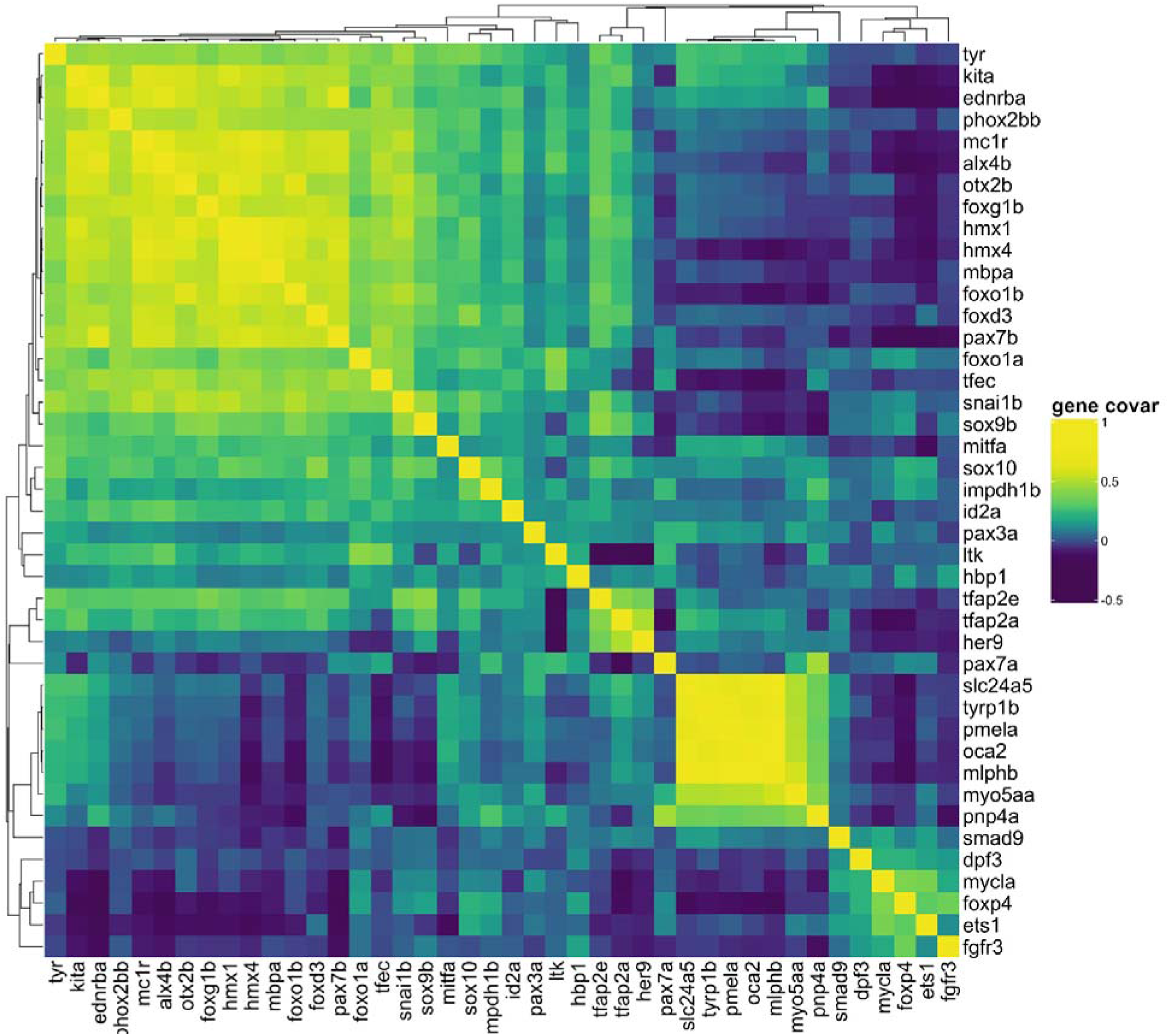
Covariance of genes within WT cells.

**Extended Data Fig. 9.**
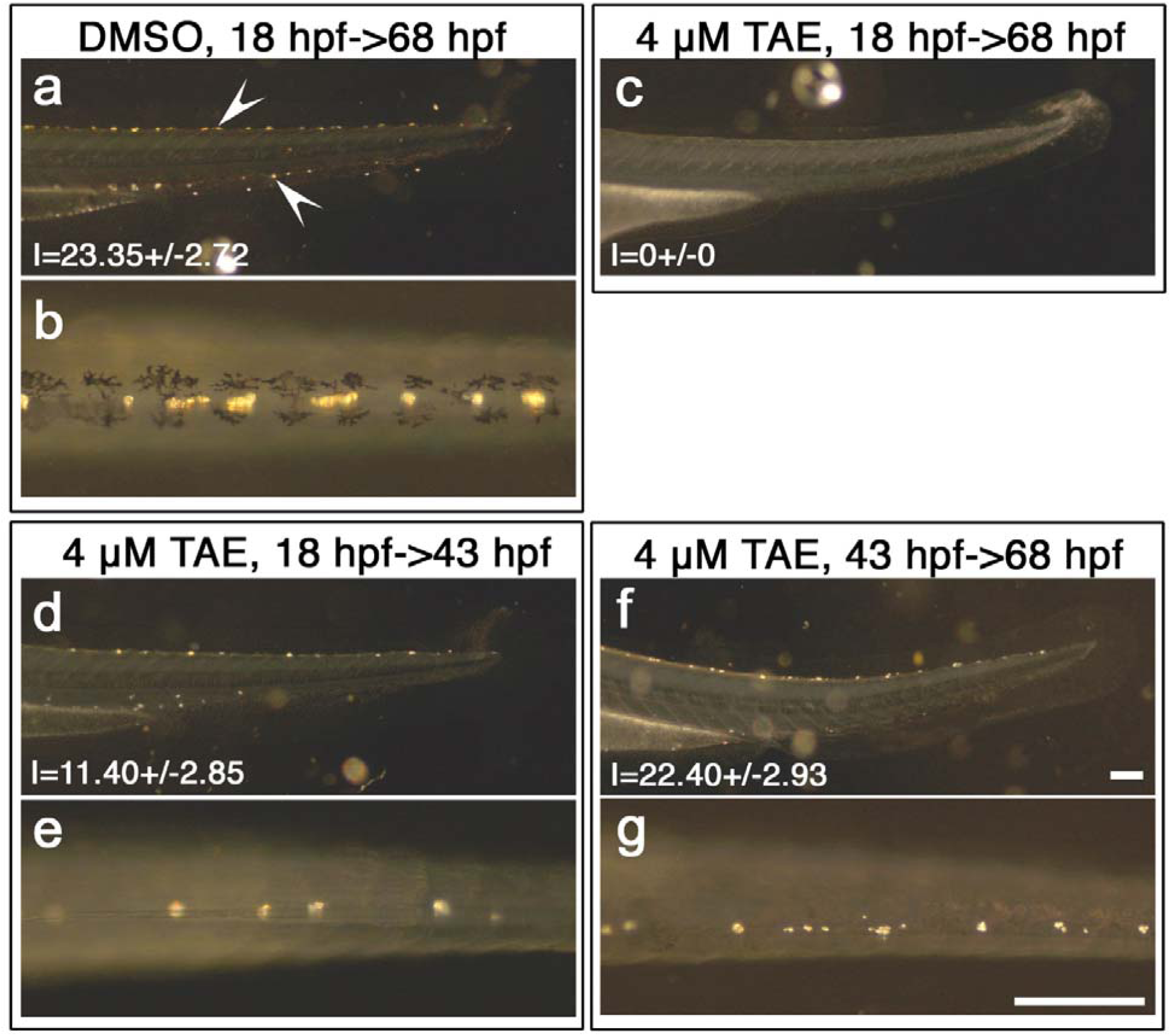
Inhibition of Ltk activity during early and late time windows results in two distinct phenotypes. Embryos were treated with DMSO (**a**,**b**) or 4 µM TAE684 (**c-g**) during defined time windows, 18 - 68 hpf (‘Control’, **a-c**), 18 - 43 hpf (‘early’, **d**, **e**) and 43 - 68 hpf (‘late’, **f**, **g**). Iridophores in dorsal and ventral stripes (arrowheads in **a**) were detected by their endogenous reflectivity. Note how ‘early’ inhibition reduces number of iridophores (specification defect), whereas late inhibition prevents their enlargement (proliferation or differentiation). (**a**, **c**, **d**, **f**) Left side views with dorsal to the top. (**b**, **e**, **g**) Dorsal views with anterior to the left. Numbers represent mean±s.d. counts of iridophores (N≥20 in each case). Scale bars, 200 µm.

**Extended Data Table 1.** Inactive version of NPM-Ltk has minimal activity to induce precocious/ectopic iridophores.

|  | No. of embryos injected | Normal morphology |  | Dead or deformed |
| --- | --- | --- | --- | --- |
|  |  | +ectopic | No additional iridophores |  |
| *Tg(sox10: NPM-Itk(DK))* | 327 (100%) | 6 (1.8%) | 240(73.4%) | 81(24.8%) |
| *Tg(sox10: NPM-Itk)* | 323 (100%) | 69 (21.4%) | 170(52.6%) | 84(26.0%) |

Notes :
“+ectopic” here includes embryos with precocious and/or ectopic iridophores. Deformed means serious defects in body axis, making it difficult to identify iridophores in normal position.

