## Supplementary Methods for "Zebrafish pigment cells develop directly from persistent highly multipotent progenitors"

**Single cell sorting and single cell data analysis**

**Supplementary file to the manuscript** Nikaido et al. Zebrafish pigment cells develop directly from persistent highly multipotent progenitors.

**Single cell sorting**

**FACS isolation of GFP+ NCCs**

Zebrafish embryos were decapitated and dissociated to a single cell suspension. Cells were excited with 488 nm laser and gated (P1) using forward and side scatter parameters to avoid debris and smallest particles. SYTOX™ Blue Dead cell stain (ThermoFisher Scientific) was added to final concentration 1 μM. Viable SYTOX™ Blue-negative; GFP-positive cells (P2) were gated using the 405 nm laser and 450/50 nm filter (DAPI channel) and 488 nm laser and 450 nm filter (GFP/FICT channel). Cells were sorted using 488 nm laser and 530/30 nm filter (GFP/FITC channel), and the 561 nm laser and 582/15 nm filter (DsRed channel). Thresholds were established by comparison with wild type fish of the same age (not shown), and a distinct population of bright GFP-positive cells (P3; Supplementary Figure 1A) was selected while bright red cells were excluded to avoid auto-fluorescent particles. Selected cells were confirmed by fluorescent microscopy (Supplementary Figure 1B).

A

**B**

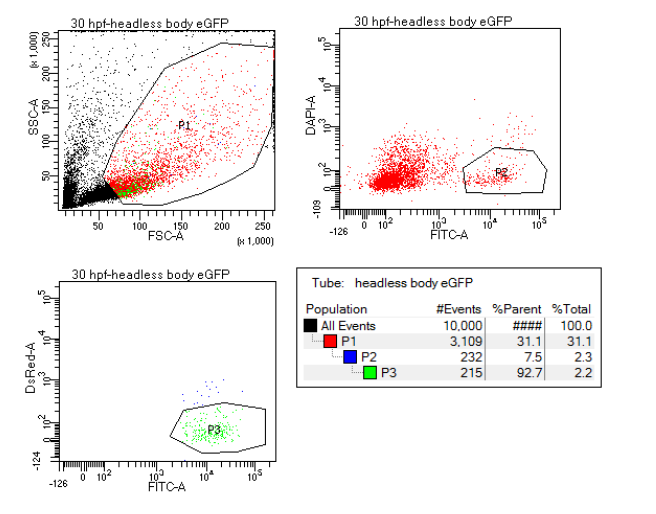

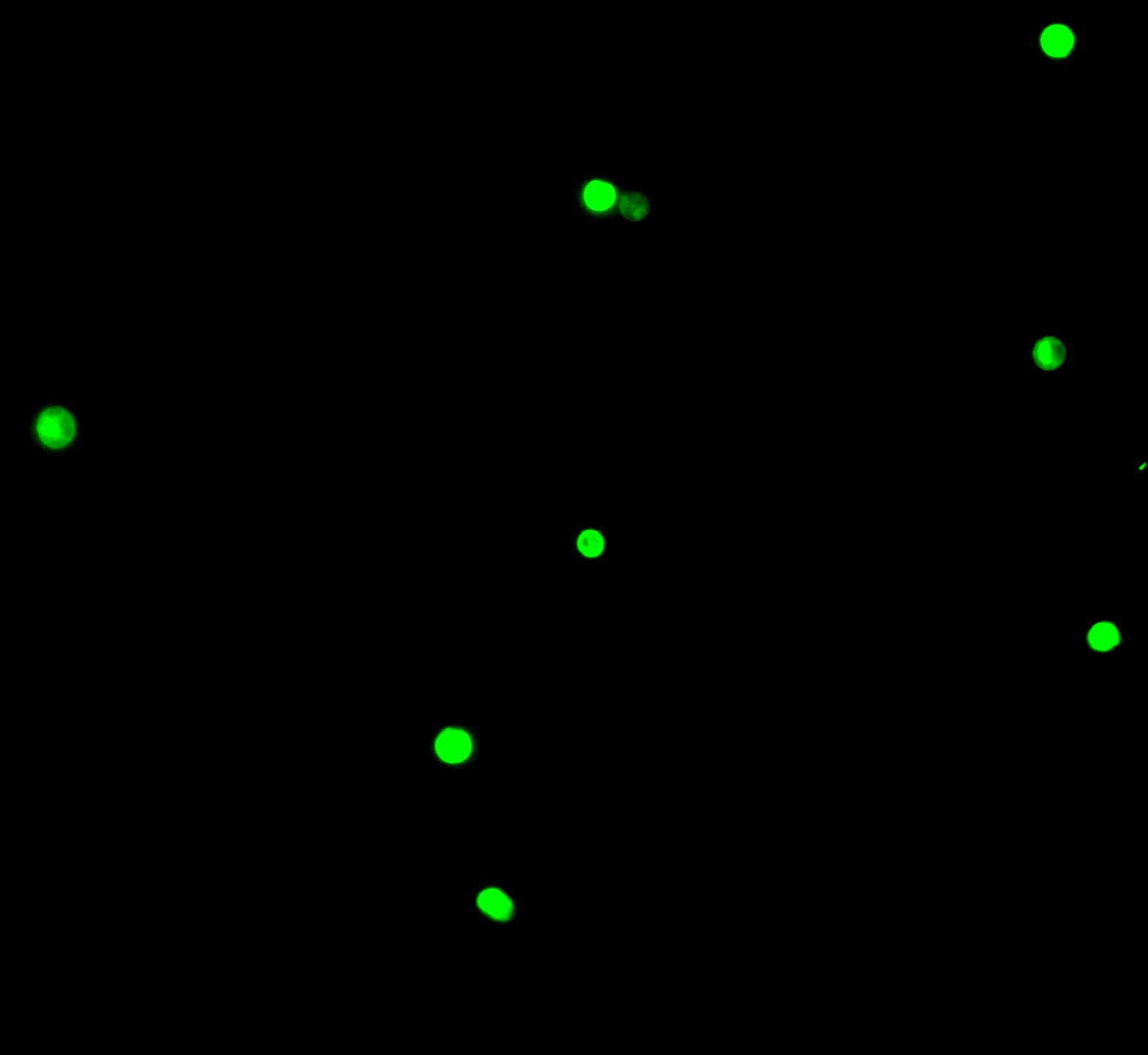

*Supplementary Figure 1 Isolation of GFP-positive (Sox10-positive) neural crest cells (36 hpf stage used as an example)*

**FACS isolation of control melanocytes and iridophores**

Zebrafish embryos (72 hpf) were decapitated and dissociated to a single cell suspension (Bi) following by centrifugation using a Percoll density gradient. The cells from the bottom of the tube (Bii) were resuspended and sorted with Aria III cell sorter using natural cell optical properties in red (DsRed) and green (FITC-A) channels. FACS plot (A) demonstrates the relative positions of melanocyte and iridophore gating. Sorted melanocytes were selected from lower right polygon (red dots) and iridophores were selected from long upper polygon (green dots). Isolated melanocytes were imaged in bright field (Biii), and iridophores were imaged using both green and red channels (shown as a merged channels black-and -white image) (Biv).

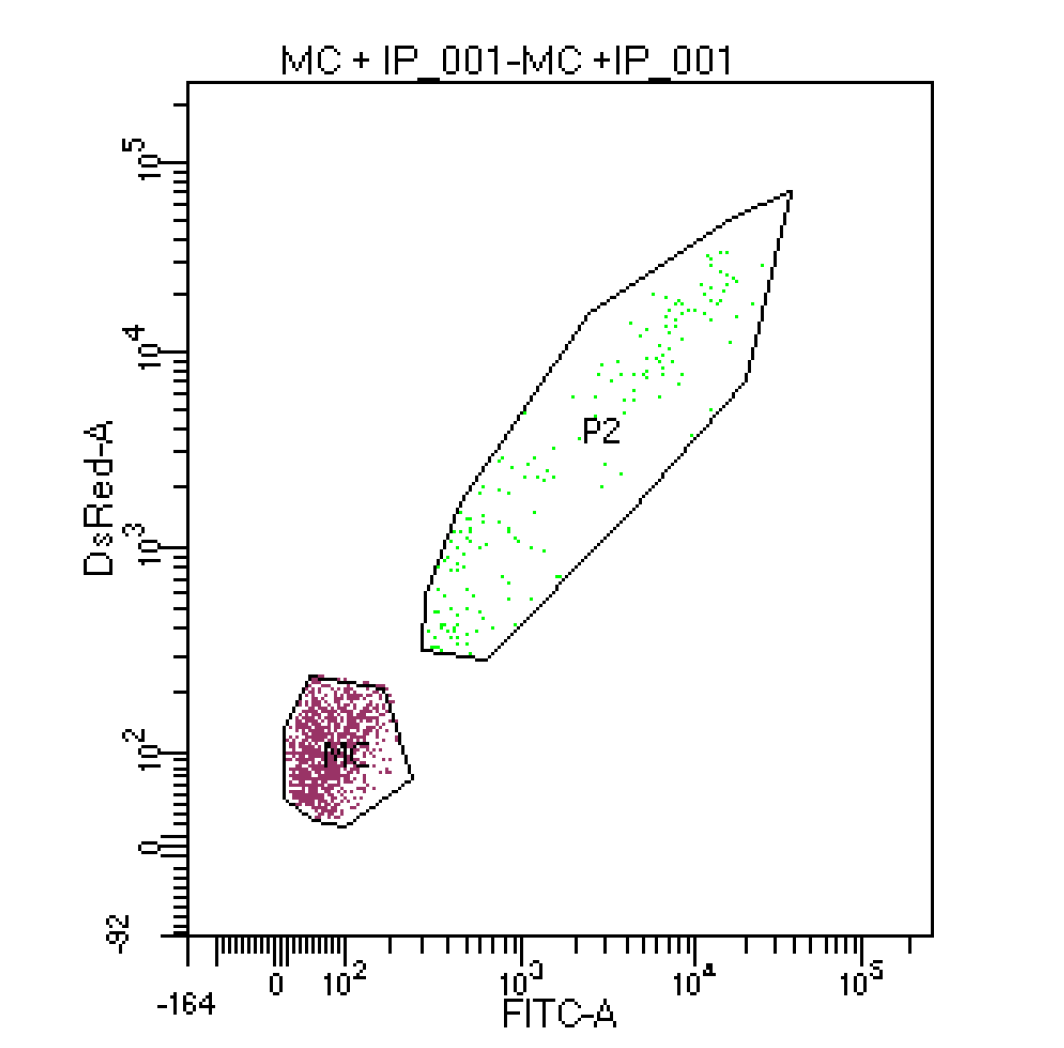

*Supplementary Figure 2* Isolation of control melanocytes and iridophores from 72 hpf wild-type zebrafish.

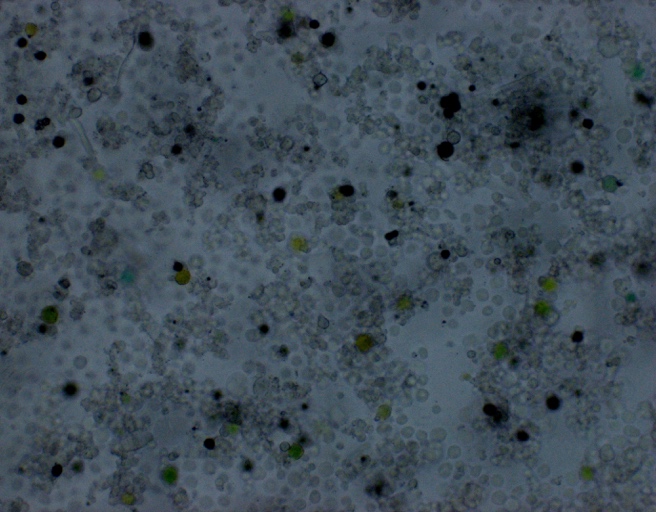

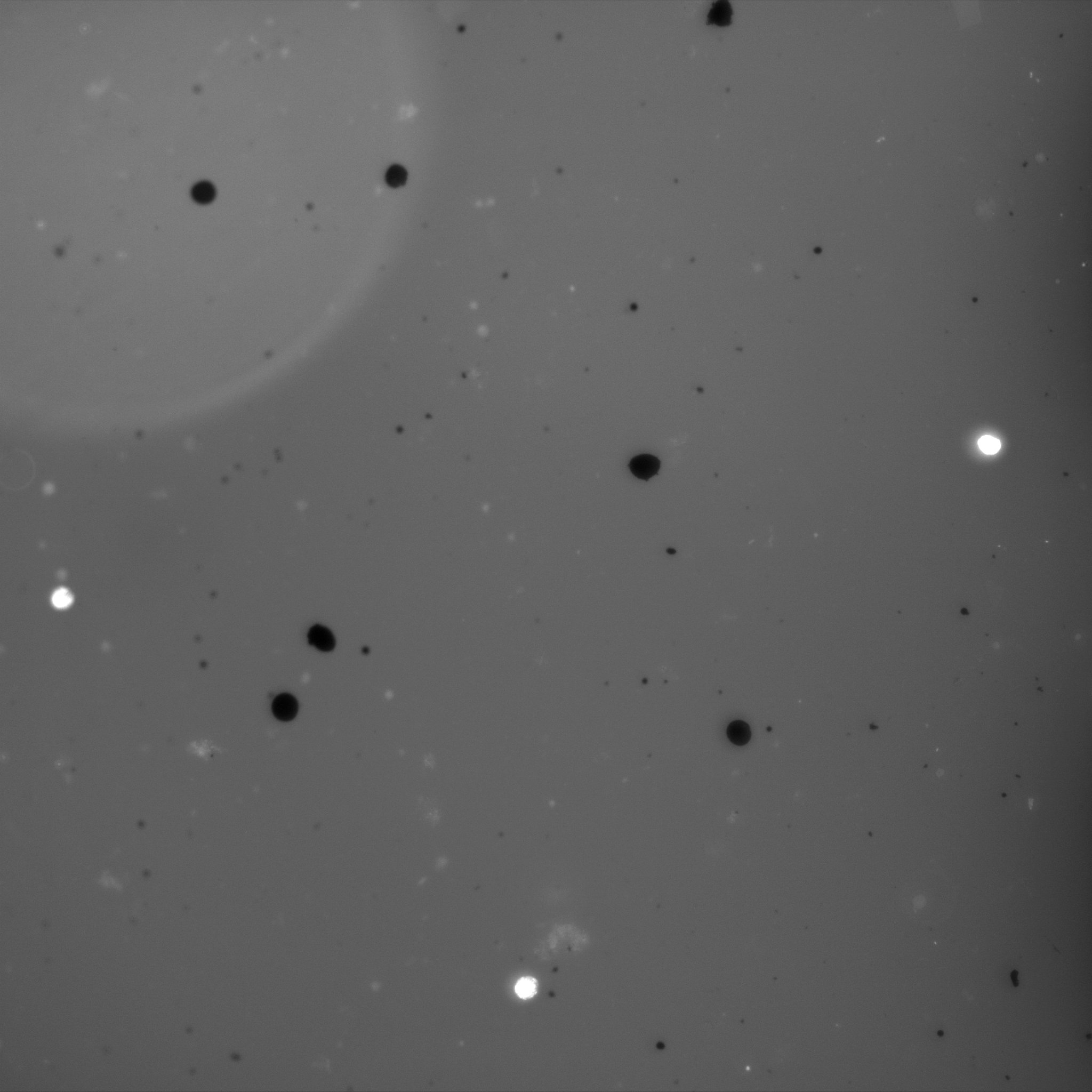

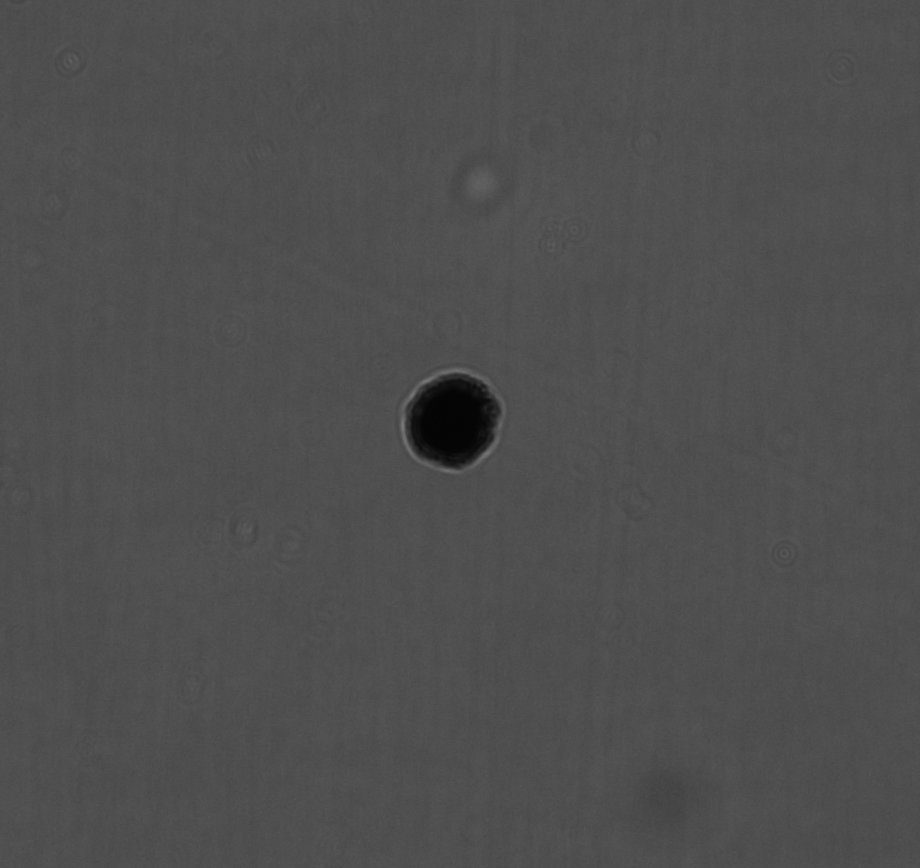

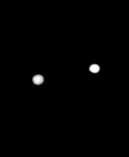

Bi

Bii

Biii

Biv

**Single cell data analysis**

**Repository:** Scripts are found in

[https://github.com/SevaVigg/NanostringDanioNCCscAnalysis/releases/tag/v1.01](https://eur01.safelinks.protection.outlook.com/?url=https%3A%2F%2Fgithub.com%2FSevaVigg%2FNanostringDanioNCCscAnalysis%2Freleases%2Ftag%2Fv1.01&data=04%7C01%7Cbssrnk%40bath.ac.uk%7C413f92f5d61a47774a8908d92fd1691a%7C377e3d224ea1422db0ad8fcc89406b9e%7C0%7C0%7C637593398724355617%7CUnknown%7CTWFpbGZsb3d8eyJWIjoiMC4wLjAwMDAiLCJQIjoiV2luMzIiLCJBTiI6Ik1haWwiLCJXVCI6Mn0%3D%7C1000&sdata=FEuU7qZz85ca%2BYyRetGLrOQG6UxafoXFqzoYQHvRfCE%3D&reserved=0)

and published at [https://doi.org/10.5281/zenodo.4953911](https://eur01.safelinks.protection.outlook.com/?url=https%3A%2F%2Fdoi.org%2F10.5281%2Fzenodo.4953911&data=04%7C01%7Cbssrnk%40bath.ac.uk%7C73ebff44efde48ec0bce08d92fd3b95a%7C377e3d224ea1422db0ad8fcc89406b9e%7C0%7C0%7C637593408661316372%7CUnknown%7CTWFpbGZsb3d8eyJWIjoiMC4wLjAwMDAiLCJQIjoiV2luMzIiLCJBTiI6Ik1haWwiLCJXVCI6Mn0%3D%7C1000&sdata=QsAxHjCpWH2PCjPPoevZrjQCVYDkyL%2BswPVrdOr%2FrwY%3D&reserved=0)

The program entry point is ProcessData.r , this script runs other scripts performing data reading, quality control, normalization, data analysis, and figure plotting.

**Nanostring nCounter® Data Analysis**

**The initial data**: The results of Nanostring transcriptome profiling are stored in 75 files. Each file (a batch of measurements) includes a header (the column names), the elements of which from five on contain the cell names, which in turn contains information about each cell, such as the experiment data, stage (hpf), and for the control cells the reference to the cell type (“IP”, “MC”, ”tail”, and “m618 sox10” for *sox10-/-* mutants). Initial experiment setup included some cells for *mitfa-/-* mutants, this experiment proved to be of poor quality and was discarded. In some files information on the cell type and hpf is repeated in the second line. Gene-related nCounter values are in rows, first four columns contain gene meta-data (the gene name, annotation identifiers, class of the probe: endogenous or control, negative or positive). Columns with cell data (the fifth onwards) contain the results of probe profiling, each column being data from one cell. Endogenous probes include external spike-in kanamycin control probe. Internal control probes include probes for 6 positive (128, 32, 8, 2, 0.5 and 0.125 fM) and 8 negative controls. We have observed that positive control probe counts were practically independent from the sum of all endogenous counts or the Kanamycin probe count. On the other hand, the counts for 6 positive controls nicely correlated with concentration of control RNA, we used the regression slope as one of the quality tests for cell data.

Data for 135 cells contained corrupted or duplicated measurements and were discarded. Initial experiment included 88 cells with *mitfa-/-* mutants, these cells proved to be of poor quality and the entire collection was discarded too. Thus, we arrived at the set of 1090 cells listed in Supplementary Table 1.

Supplementary Table 1. Initial dataset of NCC cells profiled with NanoString nCounter ®

| cell type | tails | sox10- | IP | MC | Regular WT | | | | | | | |
| --- | --- | --- | --- | --- | --- | --- | --- | --- | --- | --- | --- | --- |
| time, hpf | 24 | 30 | 72 | 72 | 18 | 21 | 24 | 30 | 36 | 48 | 60 | 72 |
| #cells | 132 | 179 | 29 | 22 | 60 | 83 | 109 | 78 | 72 | 146 | 96 | 84 |

**Data quality filtering:**

**Quality filtering of the cells.** Although NanoString nCounter proved very effective, providing quantitative expression data simultaneously for a number of genes with an acceptable rate of probe dropout measurements, even a small admixture of poor-quality data resulted in deterioration of data structure in the subsequent analysis, including poor cluster separation and irrelevant pseudotime trajectories. Thus, we kept only the cells that passed a series of rather strong filters for gene expression quality and consistency. The results of our quality control (QC) pipeline are summarised in Supplementary Figure 3.

**Our QC pipeline included the following stages**:

*Removing cells with high counts at internal negative control probes*. There are 8 negative control probes. We summed the negative probe counts and remove top 3% of the cells (Supplementary Figure 3A).

*Removing cells with poor agreement between measured and test concentration for internal positive control probes*. There are 6 positive control probes with concentrations decreasing as results of four-fold dilution (128, 32, 8, 2, 0.5 and 0.125 fM). In most of the cells positive probe counts nicely followed a similar exponential law. We used logarithms of probe concentrations as predictors and calculated the regression slope of log-counts. We discarded outlying 1% cells at each of the top and bottom (Supplementary Figure 3B).

*Removing cells with poor agreement between external spike-in (kanamycin) and internal (rpl13 housekeeping gene) control probes*. We used one external spike-in control (kanamycin) and one internal control (*rpl13* housekeeping gene). In the following analysis we used the sum of *rpl13* and *kanamycin* probes for normalization of all gene probes, so any inconsistency in the values of these two probes seriously affected normalized data consistency. We removed all the cells with disagreeing values of these probes, which often resulted from dropouts at one of the two probes. To this end we calculated regression using more stable *kanamycin* counts as predictors and *rpl13* counts as values. We removed 2.5% of all cells with residuals at the top and bottom, 5% of cells in total (Supplementary Figure 3C).

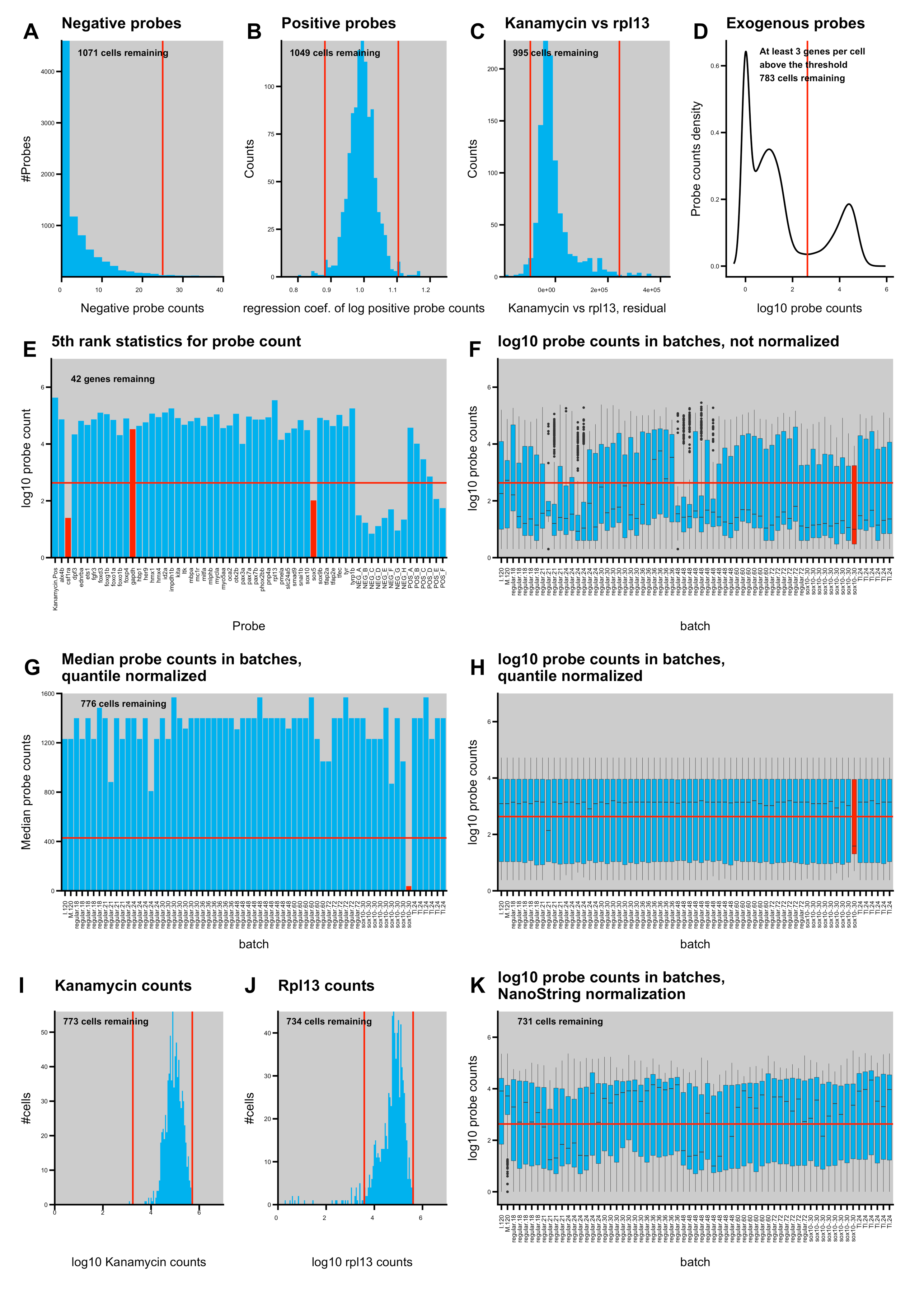

Supplementary Figure 3. Quality control and filtering pipeline of NanoString nCounter ® data. Numbers show the numbers of cells after each test. Panels: (A) Negative control probe scores. Discarded cells are to the right from the red line (top 3%); (B) Positive control scores normalized regression slopes; (C) Residuals of the regression of the external spike-in kanamycin and internal housekeeping rpl13 control probes; (D) Distribution of values of pooled probes other than spike-in kanamycin and internal housekeeping rpl13; (E) The 5^th^ rank statistics of gene expressions sums over the cells; discarded genes are shown with red bars, gapdh was initially included as an additional housekeeping internal control, but was discarded too due to its unstable expression; (F-H) distribution of probe counts in batches, (F) boxplots, no normalization, (G) medians, quantile normalization, (H) boxplots, quantile normalization; the batch marked with the red bar was discarded; (I) Distribution of spike-in kanamycin control. Discarded cells deviate from the central peak by more than the red lines; (J) Distribution of internal housekeeping rpl13 control. Discarded cells deviate from the central peak by more than the red lines; (K) Boxplots of probe counts in batches after normalization with NanoStringNorm based on the sum of kanamycin and rpl13.

*Discarding probes with poor expression of target genes in all cells.* We have observed that in a number of cells the only probes with high counts were the probes for *kanamycin* and *rpl13* controls, whereas the counts of all probes for genes of interest were comparable with negative controls. We have constructed an empirical distribution of probe values pooling values for all probes and cells (Supplementary Figure 3D). The distribution clearly had three maxima, the left hand one at zero, the central one near the average negative probe count at about 10, and the right hand one at about 25000, apparently corresponding to the actively expressed genes. We put the threshold at the trough (428 counts, shown with a red line in Supplementary Figure 1D), and kept only cells with more than three probes counting higher than this threshold. This filter proved to be the most severe, with 222 cells removed. Probably many of the removed cells are *bona fide* NCC cells adopting fates other than pigment ones and thus expressing no markers included in our panel for their attribution; note that our markers were explicitly focused on pigment cell development. If these cells were allowed to stay they generated one huge cluster, but they also affected imputation and normalization, which motivated us to exclude them at this stage.

*Removing probes with poor expression profiles.* We removed probes performing poorly in all cells, using the same threshold of 428 counts as in the previous filter. We required a probe to have a significant count at least in five cells: Two probes, *csf1ra* and *sox5* failed the test. We also manually excluded *gapdh*, which was initially included as the second housekeeping control gene but displayed a very variable expression with median less than that of many cell type specific genes (Supplementary Figure 3E).

*Removing cells from poor performing batches.* Initial data were supplied in 75 files containing data from 4 to 48 cells. The cells in the same file were measured in a single nCounter run. We considered data in different files as independent batches. Inspection of not-normalized probe counts in batches (pooled for all genes and cells) displayed large count variation (see boxplots in Supplementary Figure 3F). We analyzed distribution of probe counts in batches after quantile normalization. For small gene panels for cells of many types quantile normalization is often not the method of choice, as it magnifies expression of probes with low counts but small count ranking, e.g. in cells with a small number of actively transcribed genes included within the panel. However, in the case of cell+gene pooling in a batch this effect is much less significant, thus we performed quantile normalization in each batch. We observed that medians became remarkably well aligned with one exceptional batch for sox10 mutants (Supplementary Figure 1G,H). We removed the cells from this batch from our dataset. For quantile normalization we used normalize.quantiles()function from preprocessCore R package (Bolstad et al., 2003);

*Removing cells with a very low external control (*kanamycin*) count or internal control (*rpl13*) count*: Despite consistency test on *kanamycin* and *rpl13* expressions, there were a number of cells with very low count values for both normalization probes. As the sum of *kanamycin* and *rpl13* counts was used for normalization, simultaneous low counts of the two probes would inflate expression values. We removed 3 cells for which *kanamycin* counts were less than 0.02 of the probe median. For *rpl13* we had a number of cells with low expression and removed the cells with bottom 5% of *rpl13* probe counts (Supplementary Figure 3I,J). This ends the description of the filters we used, but a number of cells (42) were removed during normalization.

**Normalization of probe counts.** We used normalization for housekeeping genes implemented in the NanoStringNorm R package (Waggott et al., 2012). We used mean.sd for the background estimation and the sum of housekeeping gene counts (*kanamycin* and *rpl13*) for estimation of RNA content. We performed normalization in cycles, at each iteration retaining only the cells with the following conditions (as recommended by the authors of (Waggot et al., 2012): normfactors between 0.3 and 3; estimated background within 3 standard deviations from the mean background; the proportion of missing expressed endogenous probes less than 0.9 (i.e. at least 5 genes out of 42 had non-zero counts); and the estimated RNA content within 3 standard deviations from the mean. After each iteration a number of cells not complying with these conditions were removed and another round of normalization was executed until no cells were removed after normalization. For our dataset normalization stopped at the 5^th^ iteration. Some probes after normalization obtained very low values, less than one count, such values were truncated to zero. The resulting expression table was used for the imputation procedure.

**Imputation for dropouts.** Sometimes, in single cell transcriptomics a specific gene displays zero transcription activity in a cell, despite the combination of other highly expressed markers indicating that the cell belongs to a specific type, which would therefore be expected to have a high transcription activity of that specific gene. Such ‘dropouts’ are deemed to emerge from failures of reverse transcriptase initiation, but also may result from biological variability, e.g. due to transcriptional bursting (Hendriks et al., 2019). A number of software tools has been developed to correct this type of experimental error. We used drImpute software (Gong et al. 2018). As nCounter can introduce a counting error at the level of negative probe counts all elements of the expression matrix with counts less than 30 were set to zero. Then the pseudocount of 1 was added to each matrix element and log10 transformation was conducted. To exclude the possibility of a systematic influence of *sox10* mutants, the imputation was performed twice, for WT cells and for combined WT and *sox10* mutants. In total about 20% of initially zero counts have been imputed into meaningful quantitative values (2138 out of 13353).

**Single cell data analysis.**

Supplementary Table 2 contains distribution of 731 cells surviving quality filtering and normalization, which include WT, mutant and control cell types. Comparison of Supplementary Tables 1 and 2 shows that control cell types were of better quality than the WT (i.e. proportion passing quality control was higher).

Supplementary Table 2. Distribution of cells after quality filtration and normalization

| cell type | tails | sox10- | IP | MC | Regular WT | | | | | | | |
| --- | --- | --- | --- | --- | --- | --- | --- | --- | --- | --- | --- | --- |
| time, hpf | 24 | 30 | 72 | 72 | 18 | 21 | 24 | 30 | 36 | 48 | 60 | 72 |
| #cells | 108 | 135 | 25 | 19 | 46 | 37 | 54 | 53 | 65 | 43 | 80 | 66 |

After imputation the log expression values were loaded into a Seurat object (ver. 2.3.4) (Satija et al., 2015). When creating an object Seurat conducts another normalization round, thus cell datasets containing WT and WT+sox10 double mutants were uploaded to Seurat separately. Some probes were assigned low counts after imputation, thus all matrix values of less than 5 have been nullified.

**Descriptive statistics.** Seurat was used as a platform for single cell data management. We started from a principle component analysis (PCA). PCA is a linear transformation of the expression matrix thus it is likely to create fewer artifacts in cell distribution than non-linear methods such as UMAP or tSNE. We used the standard RunPCA tool implemented in Seurat suite. We have found that regular WT cells with different stages (hpf) never formed individual clouds but were grouped with different types of control cells, practically independently of their hpf. Thus, we decided to disregard the dependence of regular cells on their hpf, and introduce a single type ‘regular’. Biologically this was not unexpected since 1) it is well-known that there is a temporal gradient along the anteroposterior body axis at any individual stage, and 2) cell-type differentiation is not synchronous, but instead occurs over a broader time-window. In the PCA space (Supplementary Figure 4) the control cell types clearly lie outside the central cloud, which contains the majority of regular cells as well as most of the *sox10* mutants, as well as the majority of tail and several iridophore cells. Different control cell types deviate from the central cloud along different components, which indicates that those cells express different characteristic gene markers. Control melanocytes are strongly displaced in the second principal component (PC2), which represents the contribution of many known melanocyte gene markers (*mlphb*, *slc24a5*, *oca2*, *tyrp1b*, *pmela*), as well as some other genes (*myo5aa*, *pnp4a*, and *tyr*), which were identified as melanocyte specific genes in bulk RNA-seq or gene expression studies (Higdon et al., 2013; Petratou et al., 2018). This melanocyte cloud includes both the control melanocytes as well as some regular NCCs, which are therefore interpreted as differentiating melanocytes emerging from the natural
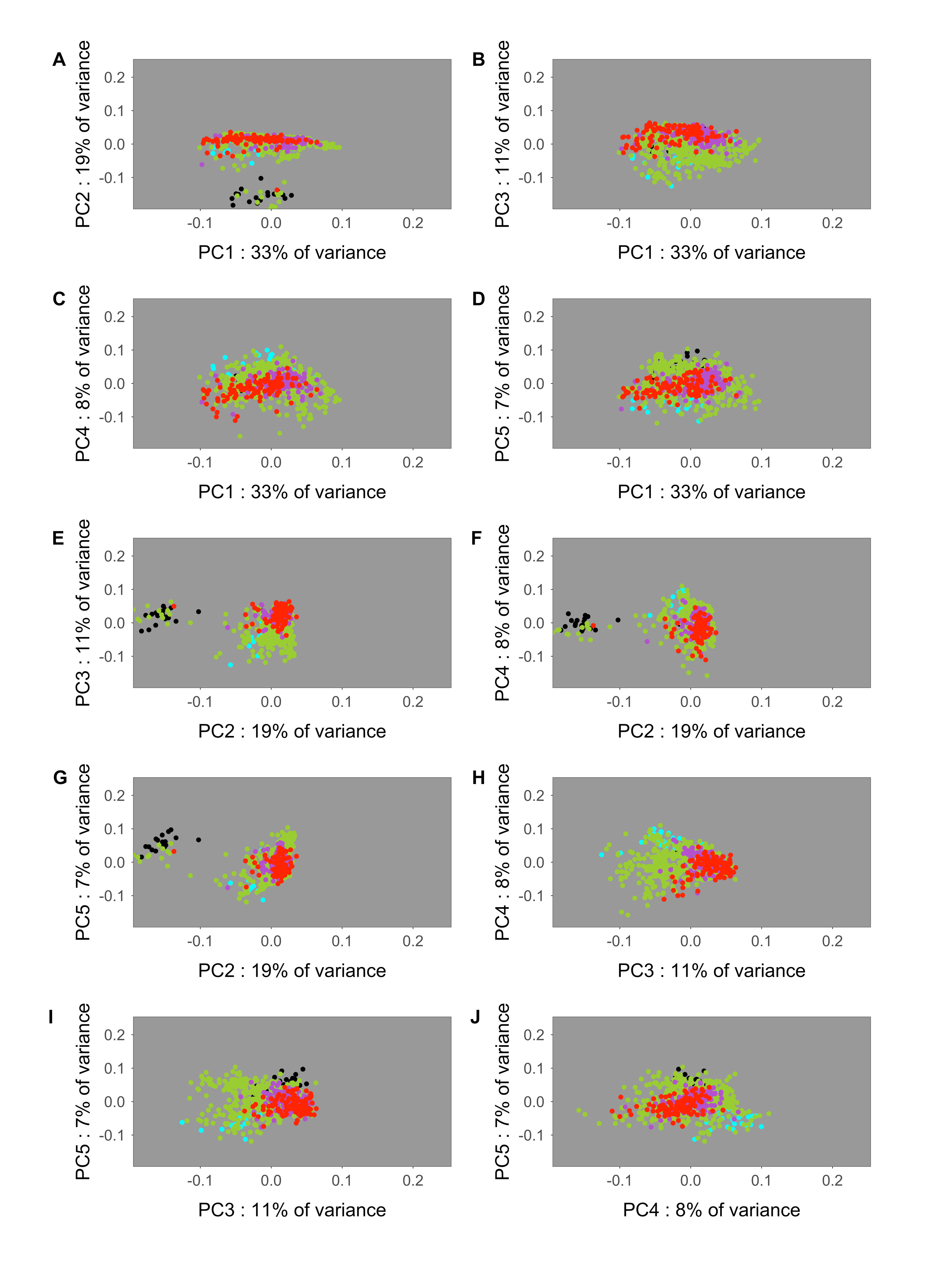

Supplementary Figure 4. Projection of cells distributions onto different pairs of principal components. Colors: (green) Regular WT cells, all hpf pooled; (red) cells taken from fish tails; (cyan) control iridophores; (black) control melanocytes; (magenta) sox10 mutants.

differentiation process. Similarly, control iridophores are found at negative values of components PC3, PC5 and positive values of PС4, in a cloud that also contains many regular WT cells, presumably differentiating iridophores. In contrast, control tail cells, used as a proxy for early NCC stages but expected to contain cells at various younger steps in NCC development, are partially offset at positive values of PC3. Interestingly, *sox10* mutant cells do not form a separate cloud but occupy a somewhat localized region within the central cloud, partly overlapping with the region occupied by control tail cells. The central positioning of *sox10* mutant cells is consistent with the previous suggestion that mutant cells are ‘trapped’ in a progenitor state (Dutton et al 2001).

**Dimension reduction maps** Even better separation of control cell types is visible after non-linear transformations using UMAP (McInnes et al., 2018) and tSNE (van der Maaten et al. 2008). We used scipy module “umap”, which we executed from R scripts using the reticulate library. For 2D UMAP visualization we used the following parameters: min_dist = 4, spread = 9, n_neighbors = 25, metric = “cosine”. To ensure that the panels in Extended Figure 1 could be readily compared we estimated parameters of the optimal UMAP transform using WT cells only, and then applied the transform to the complete data set, with *sox10* mutant cells included. For tSNE we used Rtsne R package (https://github.com/jkrijthe/Rtsne), running the optimization with the following parameters: perplexity = 20, theta = 0, eta = 500, max_iter = 50000, pca_center = FALSE, normalize = FALSE. The distance matrix for cosine distance was prepared using R package proxy.

Both UMAP and tSNE yielded topologically similar structures with clear separation of control melanocytes and, somewhat less distinctly, control iridophores (Extended Figure 1, and Supplementary Figure 8). Tail cells also grouped in a particular region of the plot.

**DotPlots**. The DotPlot tool implemented in Seurat has expression coloring normalized independently for different genes. In the case where there are small numbers of cell types this can be visually misleading, e.g. in Extended Figure 1c expression of many weakly-expressed genes (e.g. *sox10* mutant cells) will have red coloring for the sample with the highest value (contI in the case of sox10), despite a rather modest absolute count value in this cell type, because counts in other cell types are even lower. To control this problem, we have designed our version of DotPlot (dotPlotBalanced.r) which uses a coloring scheme evaluated from expression of all genes in all cells of the panel. Our implementation is based upon ggplot2. Dotplots in Supplementary Figures 3 and 4 are colored according to this scheme.

**Heatmaps** We used the ComplexHeatmap R package (Gu et al. 2016) to construct heatmaps. We used the viridis color map, for aesthetic reasons, but also to assist those with impaired colour vision. When clustering rows or column, we used cosine distance implemented in R package proxy.

**Clustering** We used the standard Seurat software FindClusters to cluster cells by their probe count profiles. This software uses the shared nearest neighbor tree and the Waltman-van Eck algorithm to identify communities. The neighbors were identified using Euclidean metrics in the UMAP image space. We fixed the value of k.param to be 15 and then conducted enumerative testing of all parameters combinations including values of UMAP target dimensions between 2 and 8, UMAP min_dist between 1 and 4, and cluster resolution between 0.5 and 6. Each round was replicated three times. To select the optimal clustering, we profited from the availability of iridophores and melanocytes as control cell types. We used these control cell types to control for gene expression variation between cells from a single cell type. Importantly, some cells naturally develop into the same cell type as these controls, and reassuringly obtain a very similar looking gene expression profile (see Extended Figure 1). If too high a value was selected for cluster resolution this would result in splitting of clusters, with control cells distributing between several clusters. Conversely, too low a cluster resolution would result in merging of several clusters with control cell type cells diluted by cells from other cell types and reduced regular cell similarity in the cluster. We designed a heuristic objective function, which has an optimum for cluster containing many cells with large proportion of target control cell type cells

$$qual=\frac{1+N_{irido}}{\sqrt{1+N_{clust\_irido}}}+\frac{1+N_{melo}}{\sqrt{1+N_{clust\_melo}}}$$

Here $N_{clust\_irido}$ , $N_{clust\_melo}$ are the total number of cells in the target cluster, whereas $N_{irido}$ , $N_{melo}$ are the number of cells of the target cell type in the target cluster. If a cluster did not contain target cells its quality was set to zero. The optimal clustering was selected as that with the maximal objective function value over all possible parameter combinations and replicates. The optimal clustering was remarkably well reproduced in different rounds of clustering (this was not the case in our initial tests, when quality filtering was less stringent) and consisted of 10 clusters. The melanocyte target cluster contained all 19 control melanocytes in a cluster of 37 cells whereas the iridophore target cluster contained 20 out of 25 iridophores in a cluster of 49 cells. We described this clustering as ‘the coarse grain clustering’ since some clusters had a similar gene expression profiled and were merged in the next stage (Supplementary Figures 5,6) .

We assigned specific cell types to clusters using either a high proportion of the control type cells, or dominant expression of particular markers. Control type cells were used to assign melanocytes and iridophore clusters. A small cluster containing 22 cells and expressing *pax7* genes *a* and *b* was identified as xanthophores. The cluster containing the greatest proportion of cells from tails were identified as the earliest in the set and were named early highly multipotent progenitors (eHMP). This cluster contained cells expressing the early markers like *sox9b* and *snai1b,* and an eHMP specific marker *tfap2e*, but in fact its cells had a broad expression, showing high counts for the majority of genes in the panel. Many of those genes are considered as early fate decision effectors, e.g. *phox2bb*, *kita*, *mitfa*, *pax7b*. Another cluster of cells with a broad gene expression pattern did not express *tfap2e*, had noticeably lower counts for *sox9b*, but had a high count of *ltk* (essentially absent in eHMP cells), and prominent expression of *foxo1a.* We named this cluster ‘ late highly multipotent progenitors’ (ltHMP).

Nevertheless, several clusters of this optimal clustering had similar sets of characteristic markers, apparently indicating similar or the same cell types. To identify only characteristic cell types, we merged similar clusters using the Seurat ValidateClusters procedure. ValidateClusters tests all pairs of clusters and uses a modification of the support vector machine classifier algorithm to identify if the pair of clusters have the same top marker genes. We performed validation in the principle component space and used the following parameters: number of principal components = 8, min.connectivity = 0.005, top.genes = 4. ValideateClusters is a greedy procedure; to avoid dependence on the order of randomly selected pairs we repeated the cluster merging procedure increasing the accuracy cutoff threshold stepwise from 0.75 by 0.01 until the control clusters (those melanocytes or iridophores) start to merge with other clusters. The first merging occurred at 0.92 with iridophores merging to xanthophores, and as a result we obtained 7 validated clusters.

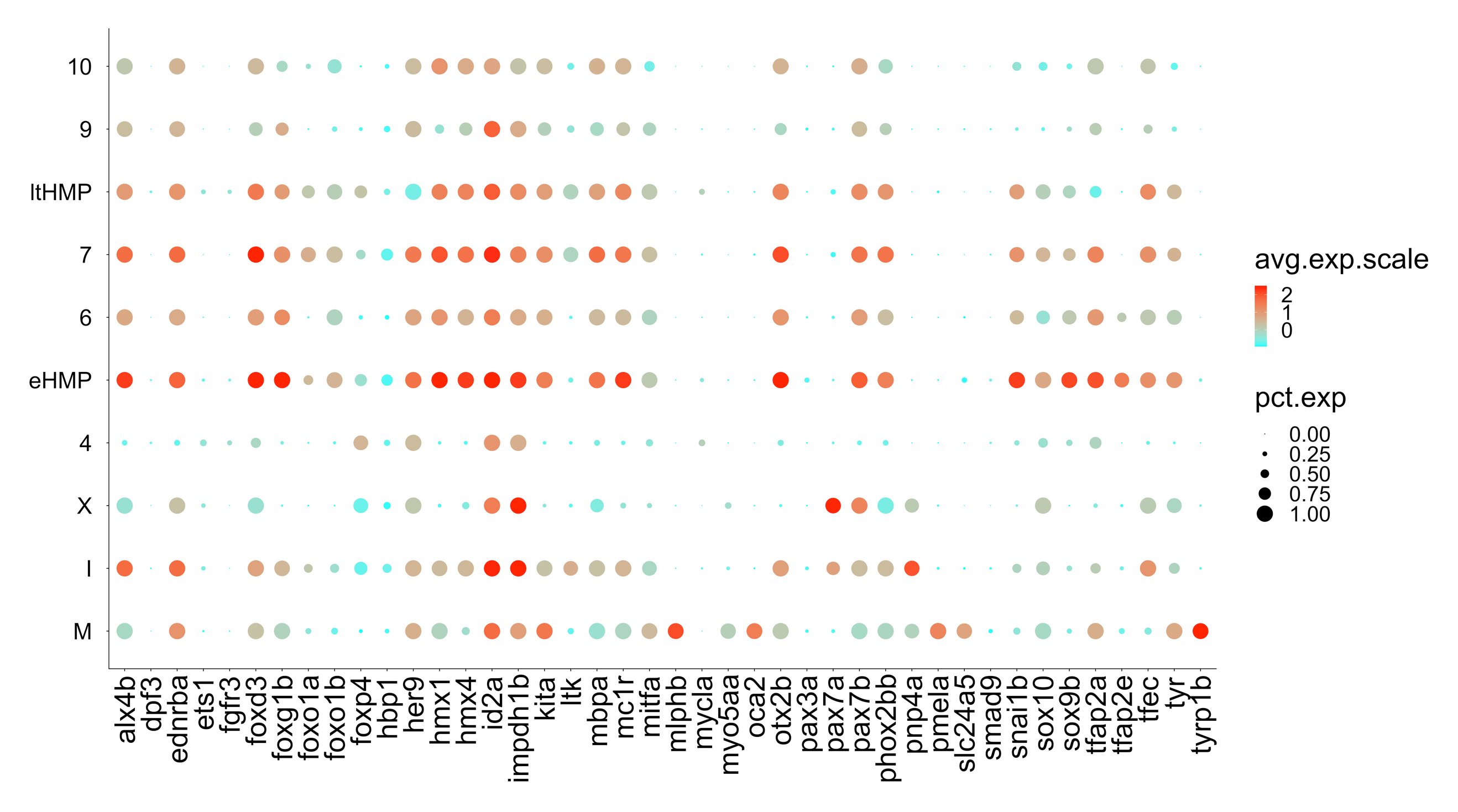

Supplementary Figure 5. Probe count profiles of genes, expressed in the coarse grain clustering. . Cluster labels (M) melanocytes; (I) iridophores; (X) xanthophores; (ltHMP) late highly multipotent progenitors; (eHMP) early highly multipotent progenitors. Clusters labeled with numbers could not be given more precise names with the markers available in the panel.

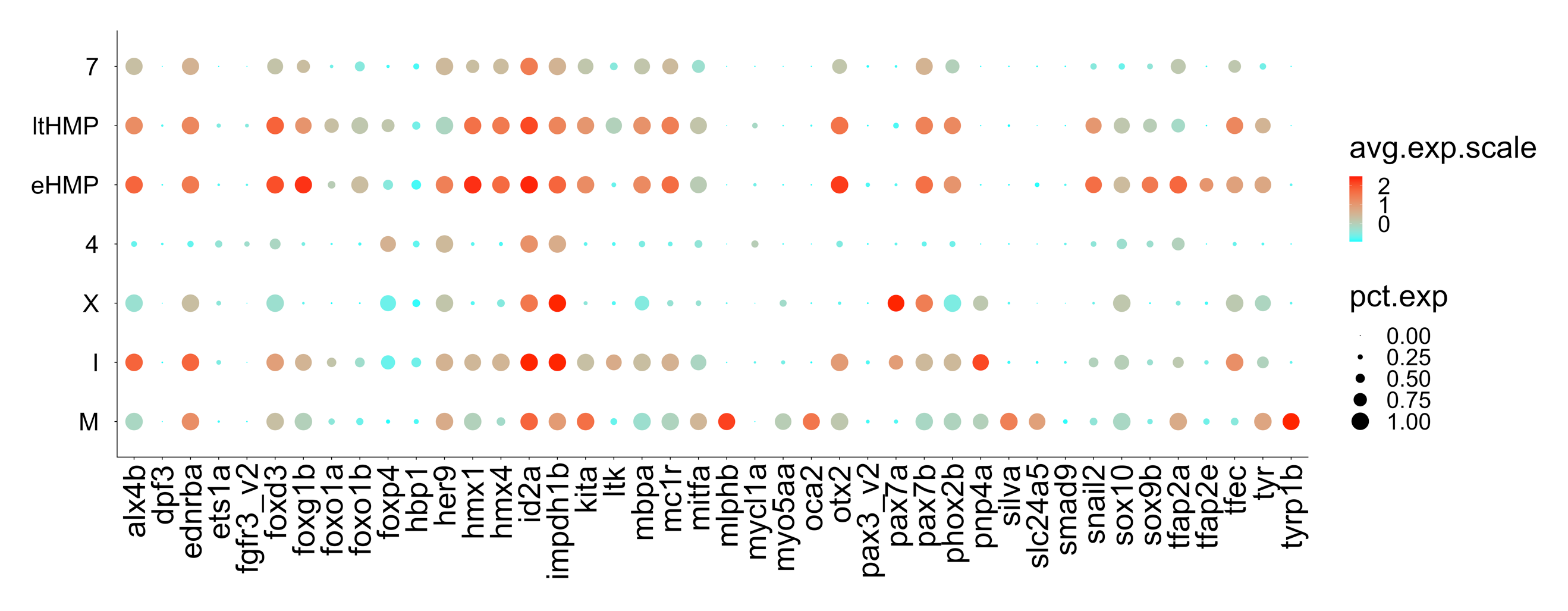

Supplementary Figure 6. Probe count profiles of the optimal clustering after cluster validation and merging of clusters with similar counts of 4 top genes in each of the 8 top principal components. Cluster labels (M) melanocytes; (I) iridophores; (X) xanthophores; (ltHMP) late highly multipotent progenitors; (eHMP) early highly multipotent progenitors. Clusters labeled with numbers could not be given more precise names with the markers available in the panel. Clusters are renumbered after cluster validation, thus clusters 4 and 7 are different from those in Supplementary Figure 3. During validation clusters 9 and 10 from Supplementary Figure 3 (SF3) merge together into cluster 7 in this Figure, cluster 7 from SF3 merges with ltHMP, and cluster 6 from SF3 merges with eHMP,

We used the BuildClusterTree tool in Seurat to construct a tree of the validated clusters. The tree is represented in Supplementary Figure 7.

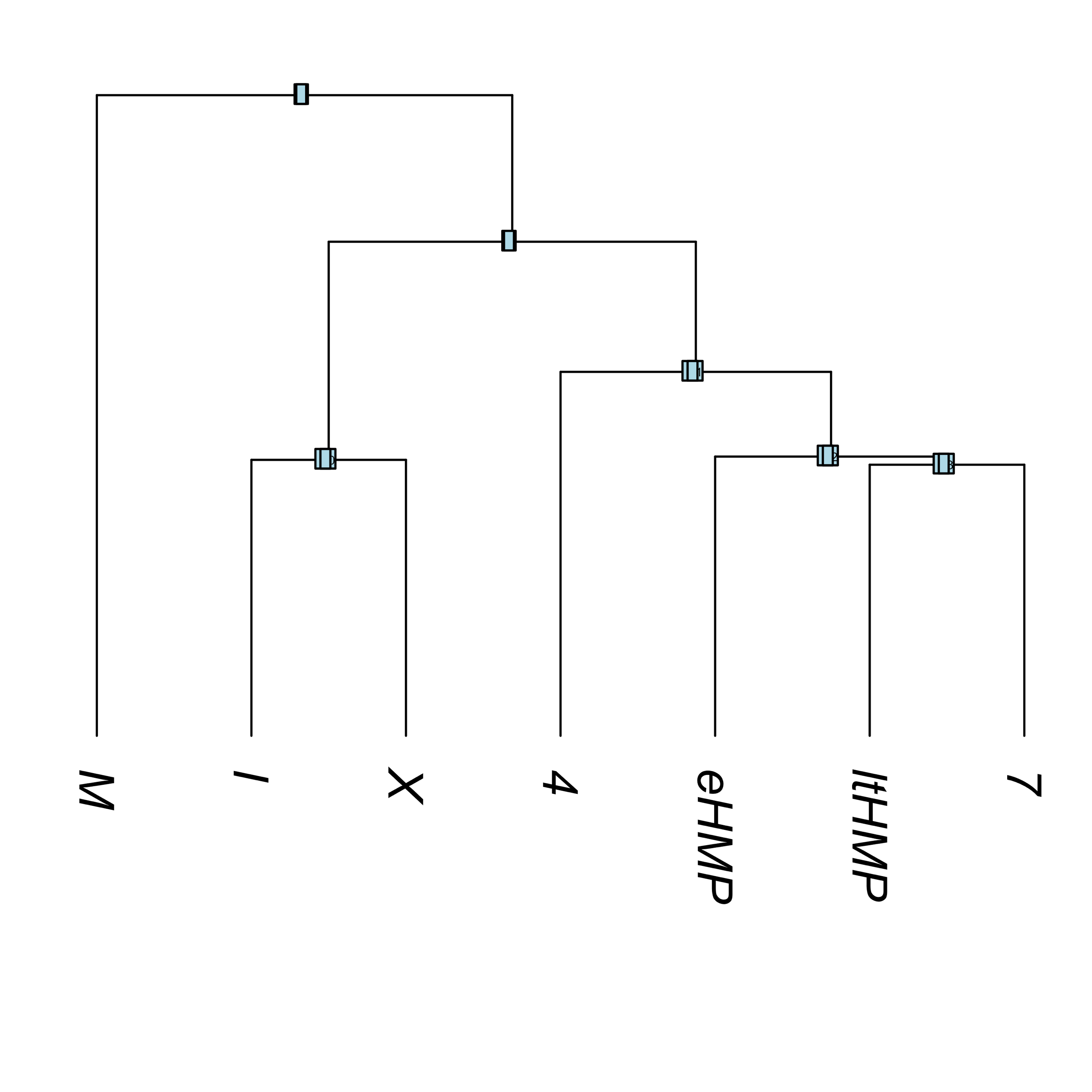

Supplementary Figure 7. Similarity tree of validated clusters of cells profiled with Nanostring panel of 42 genes. Cluster labels (M) melanocytes; (I) iridophores; (X) xanthophores; (ltHMP) late highly multipotent progenitors; (eHMP) early highly multipotent progenitors. Clusters labeled with numbers are difficult to interpret further with the number of markers available in the panel. Melanocytes are outliers due to the large number of melanocyte-specific markers in the panel.

Validated clusters identified by SNN occupy clearly delineated basins both in the 2D UMAP and 2D tSNE maps; such basin separation, especially in the tSNE plot (recall that this algorithm was not used for cluster identification), serves as an additional sanity check (Supplementary Figure 8).

| 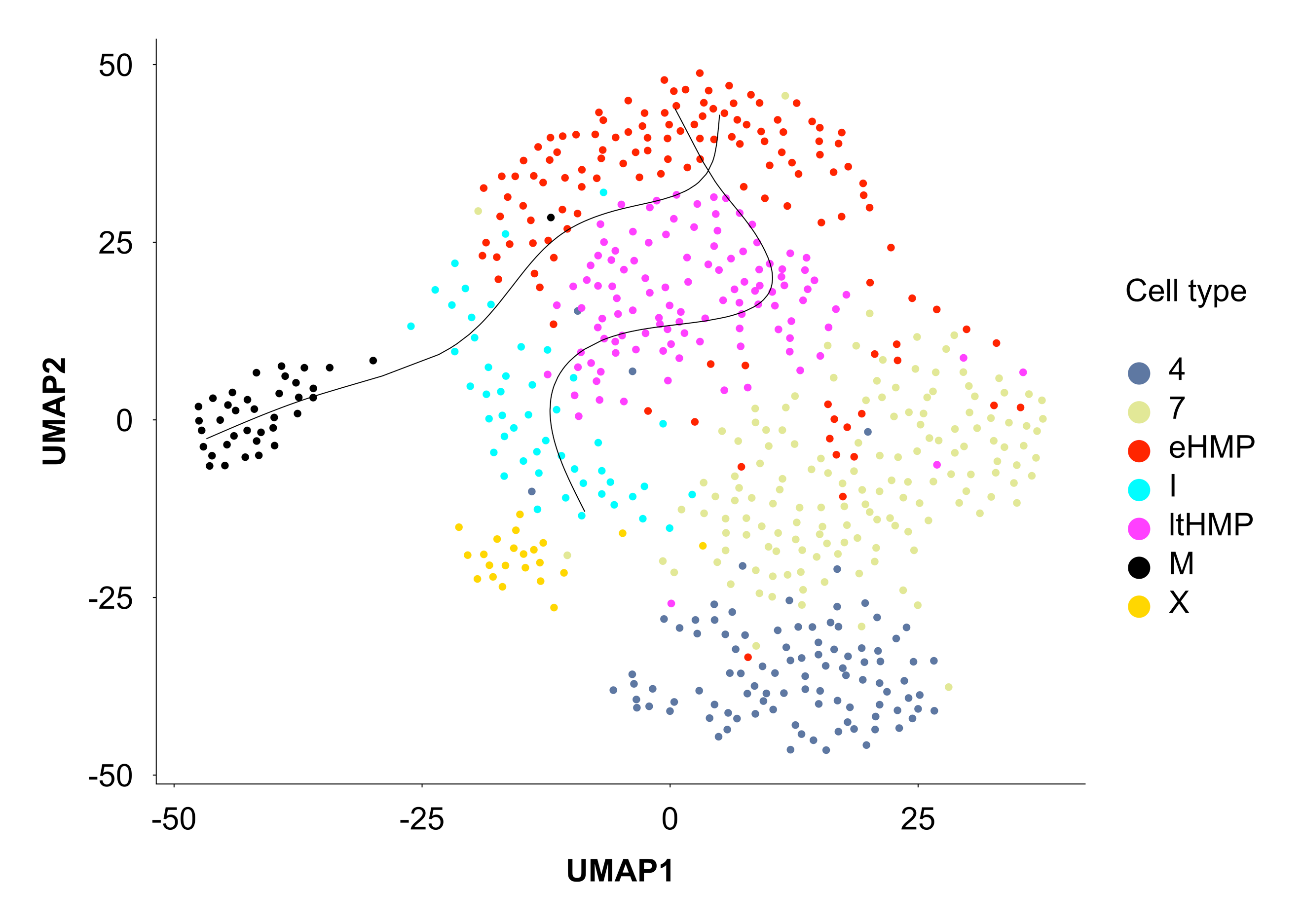 | 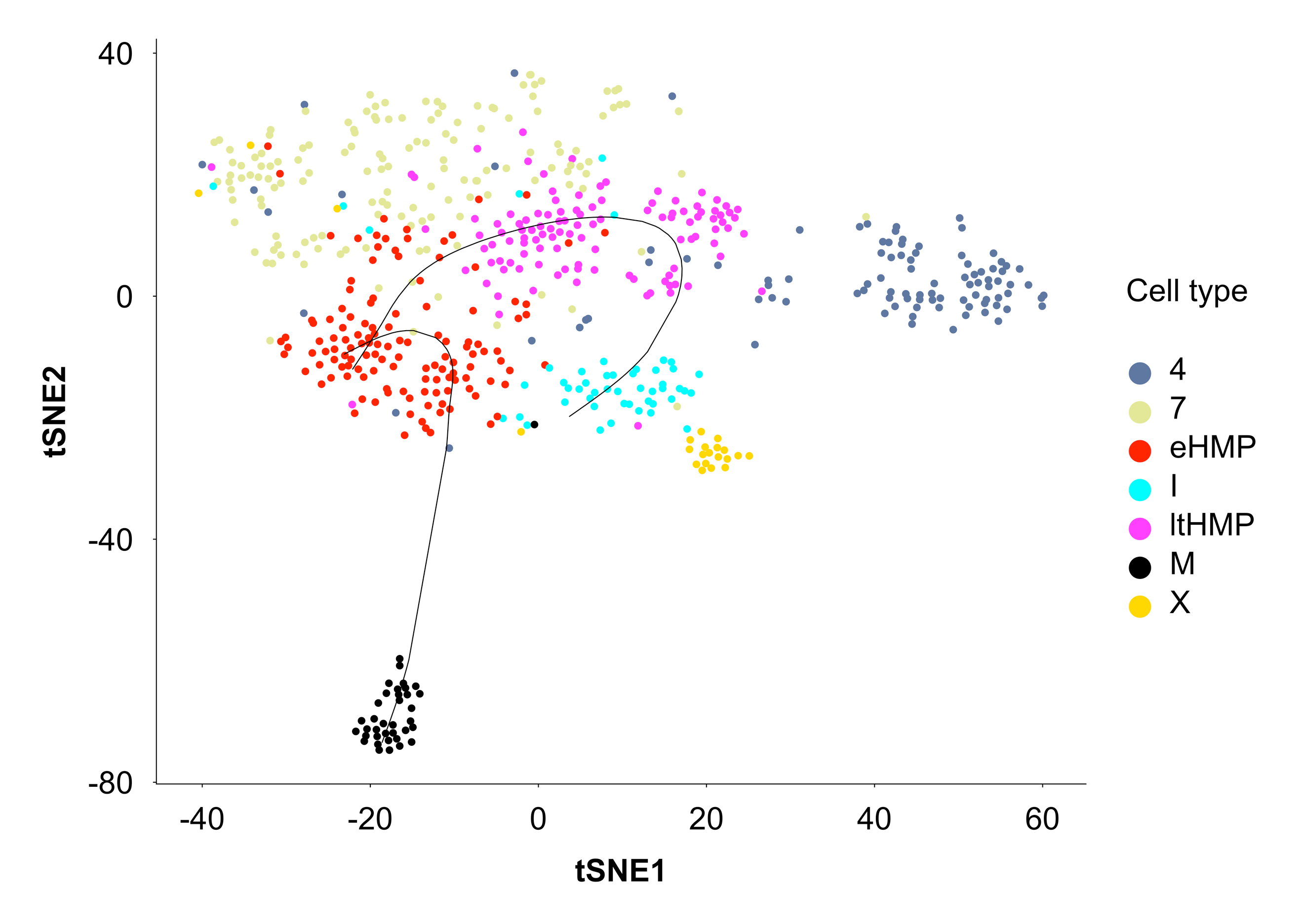 |
| --- | --- |

Supplementary Figure 8. Cluster visualization and pseudotime lineages in 2D UMAP and tSNE maps. Pseudotime lineages are constructed in 42D gene expression space by slingshot software.

**Violin plots** We used the standard Seurat tool VlnPlot. Violin plots for the clusters are given in Extended Data Figure 2.

**Feature plots** We used the standard scheme implemented in the Seurat tool FeaturePlot. In all cases a 2D UMAP visualization plane was used. We used the bottom cutoff on log10 gene expression (parameter FeaturePlot minCutoff = 3). Feature plots are represented in the Extended Data Figure 3.

**Pseudotime ordering** Program slingshot (ver. 1.0.0) (Street et al., 2018) was used to create trajectories (principle curves) in the 42-dimensional space of gene expressions. Two slingshot objects were created, one in the 42D space of gene expressions, the other in the 2D space of the UMAP image. The first was used for heatmaps (see Extended Figure 5), the second for trajectory visualization (as in Supplementary Figure 8). In both cases we selected eHMP as the starting cluster and “I” and “M” as terminal clusters of two lineages. We used the following parameters: extend = "n", reassign = TRUE, stretch = 0, thresh = 0.005, shrink = 0.4, dist.fun = “cosine”. Cosine distance between clusters is not included in standard software and was implemented from scratch as

$$D=\sqrt{2\left( 1-\frac{\sum_{i in genes} \bar{A_{i}}\bar{B_{i}}}{\sqrt{\left( \sum_{i in genes} \bar{A}_{i}^{2} \right)\left( \sum_{i in genes} \bar{B}_{i}^{2} \right)}} \right)}$$

where $\bar{A_{i}}$ and $\bar{B_{i}}$ are the average of log-expression of gene *i* over all cells in clusters A and B. Here we profited from an undocumented option in *slingshot* that allows the software to accept distance functions in the form $f(X, w_{1}, w_{2})$, where *X* is the gene expression matrix and $w_{1}$ and $w_{2}$ are index vectors whose entries are 1 if a cell is included in the cluster and 0 if not.

This set of parameters allowed us to start trajectories in different regions of the starting point cluster (owing to the non-zero value of the shrink parameter), with the ends of trajectories not protruding much from the terminal cluster cloud (owing to combination of extend and stretch values).

Cells for heatmaps in Extended Data Figure 5 were obtained from the slingshot object constructed in 42D using a threshold of 0.95 to collect cells ( curveOrd[ curveWt > 0.95, ] of the slingshotObject@curves ).

**Analysis of Data Obtained by Single-Cell Gene Expression Profiling using TaqMan® Gene Expression Assays**

**Initial data:** The results of single cell TaqMan RT-PCR profiling were supplied in a table csv file (semicolon separated) and contained results of profiling of expression of 13 marker genes, the external control (*kanamycin*) and the internal housekeeping control (*rpl13*) in 159 cells. The list of genes was different from that used for Nanostring analysis: {***elavl3***, *ltk*, *mbpa*, *mitfa*, ***neurog1***, *pax7b*, *phox2bb*, *pnp4a*, *snai1b*, *sox10*, *sox9b*, *tyrp1b*, ***xdh***}, here genes not present in the panel used in Nanostring profiling are shown in bold. All TaqMan profiling was performed for cells at 30 hpf. The elements of the table contained either decimal numbers showing the number of the cycles needed to obtain the threshold signal, or the word “Undetermined” used if no signal were obtained in 40 cycles. Some elements with failed experiments contained NA symbol.

**Data processing and filtering:** First values of all elements marked as “Undetermined” were set equal to 40, the maximal number of the cycles. Then log10-expression values were calculated using the formula:

$$Expression=\left( 40-nCycles \right)*{log}_{10}(2)$$

The variance of expression values of internal housekeeping control gene *rpl13* proved to be 25-fold more variable than those for *kanamycin* (2.26 vs 0.092), thus we decided not to use *rpl13* values for normalization for RNA quantity. On the other hand, the stability of measured concentrations of *kanamycin* spike-in control were satisfactory, thus we decided to use normalization only for *kanamycin* concentration. To this end we first removed 5% of samples with the highest *kanamycin* expressions (to get rid of the outliers) and then subtracted *kanamycin* values from all gene expression values for the corresponding cell. Since the data in the table contains logarithmic values, this operation corresponds to division of non-logarithmic expression values by *kanamycin* expression. A small number of elements in the data matrix with failed experiments were marked as <NA>, we set such gene expression to zero.

*Removing the samples for small values for all genes.* Some samples had a very low normalized expression for all genes. We computed the total gene expression for samples and dropped the bottom 5% of samples.

*Transition to the relative units.* The resulting logarithmic data were negative and rather large by their absolute value. To make them more suitable for interpretation we transitioned to relative units by computing the minimal matrix element of the expression matrix and subtracting it from all matrix elements.

*Removing matrix elements with very small gene expression.* In relative units a large number of genes had expression values very close to 0 (see Supplementary Figure 9). We have found the trough and set the threshold to discard the low scoring probes in the left maximum.

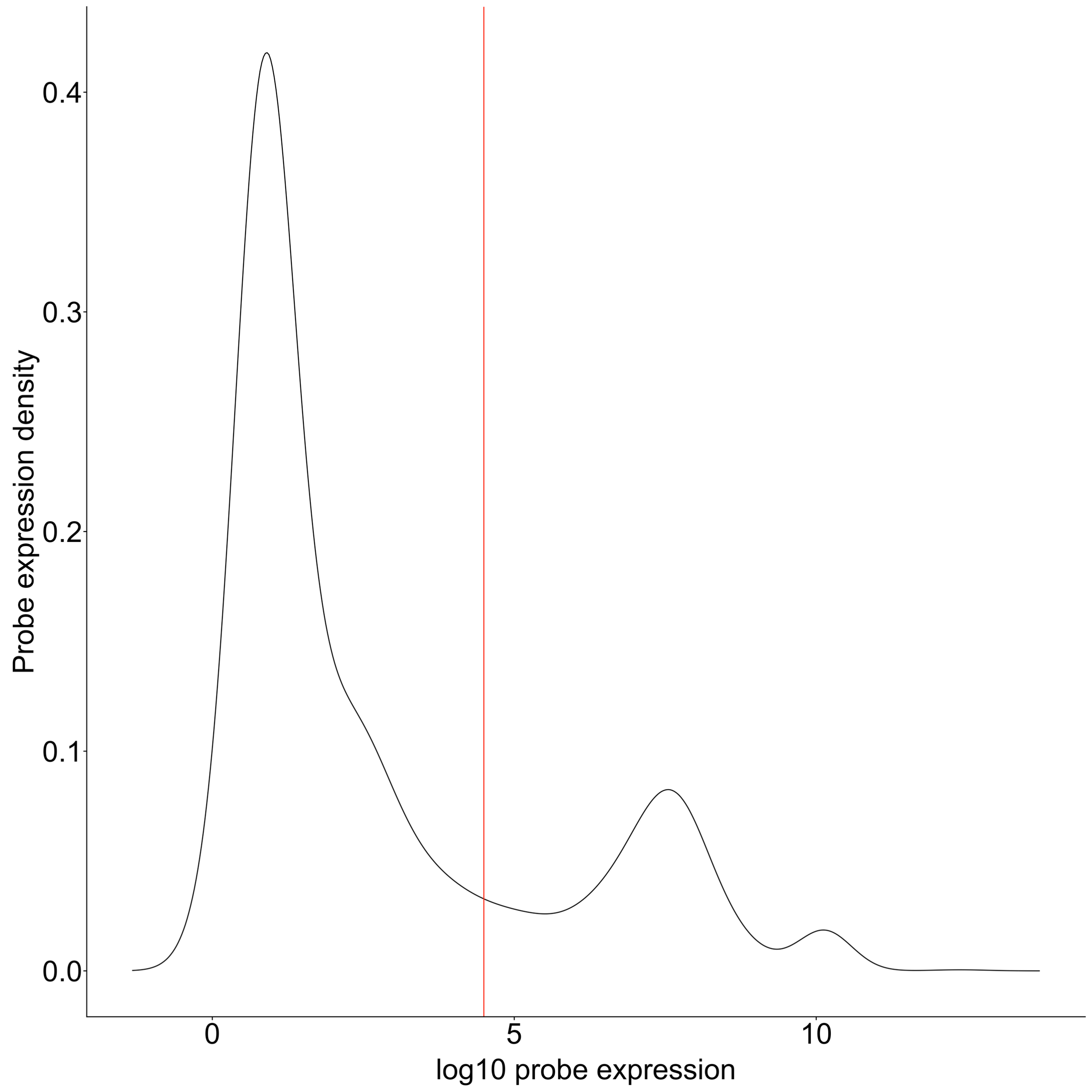

Supplementary Figure 9. Distribution of TaqMan expression assay values. Probes to the left of the red line threshold were set to zero.

The surviving probes were used to construct the Seurat object.

TaqMan data were used to build heatmaps of gene expression in samples. Supplementary Figure10 contains the complete heatmap of all TaqMan assay probes.

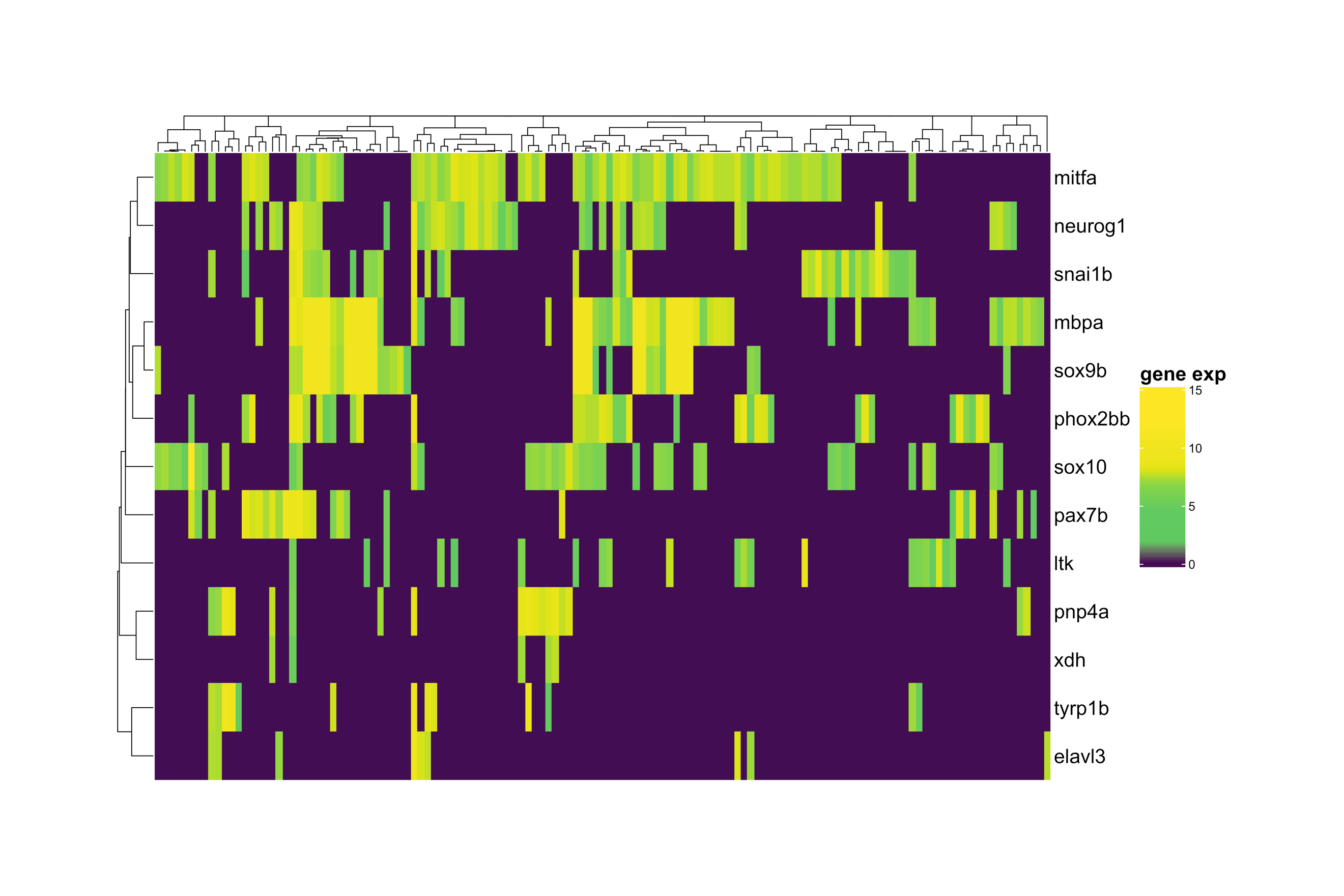

Supplementary Figure 10. Heatmap of TaqMan probes. Clustering trees for both rows and columns are built using cosine distance in the initial space of probe log-expressions in relative units recalculated from cycle numbers.

A heatmap of probes with positive *ltk* (22 probes in total) shows that *ltk* is indeed co-expressed with different genes (Supplementary Figure 11).

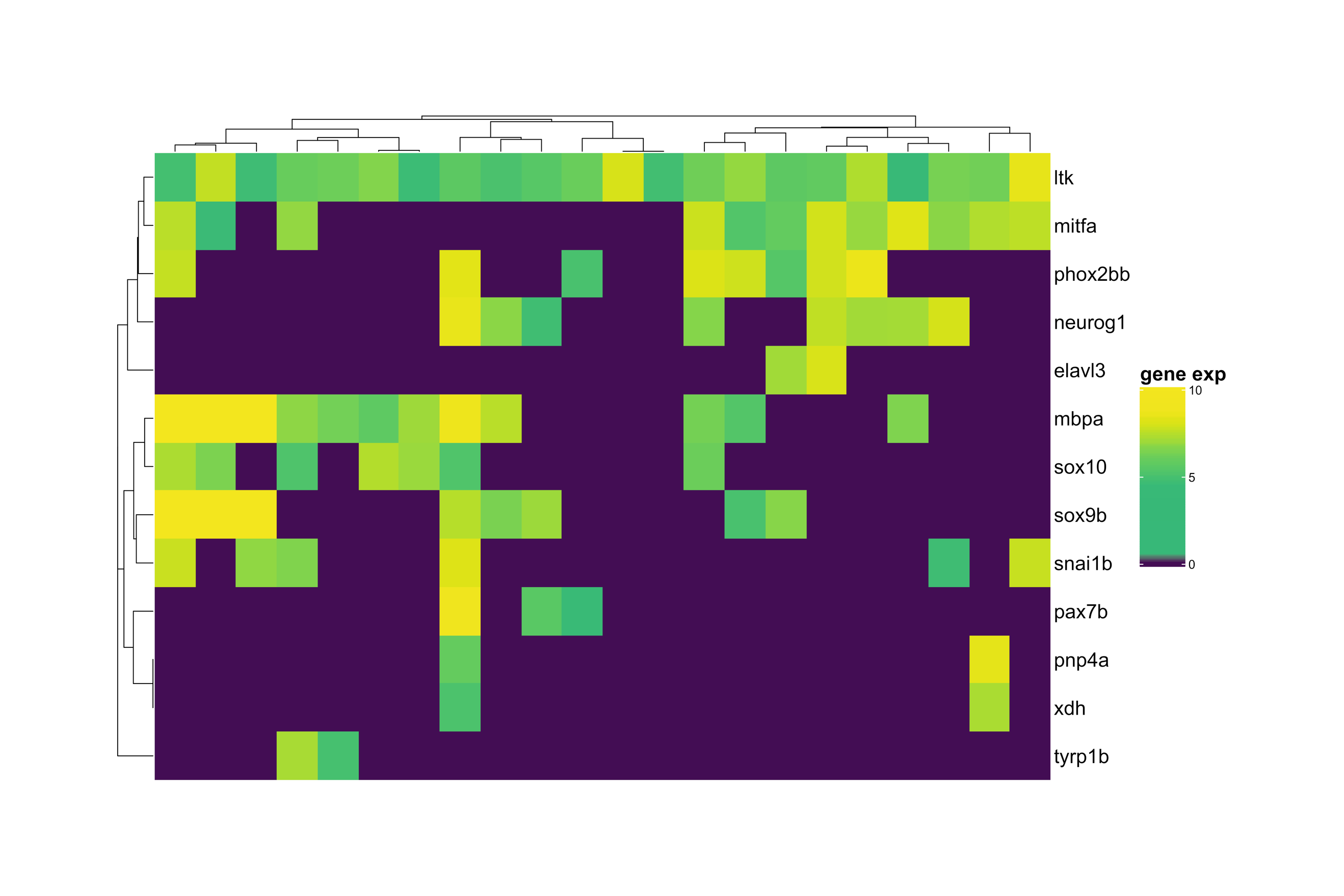

Supplementary Figure 11. TaqMan assay of samples with positive ltk expression. Note the diversity of genes co-expressing with ltk.
