## Supplementary Information for "Zebrafish pigment cells develop directly from persistent highly multipotent progenitors"

**Supplementary Methods**

**Supplementary Table 1**

List of neural crest-related genes assessed by NanoString. In addition, we assessed the housekeeping gene *ribosomal protein L13* (*rpl13),* and also *glyceraldehyde‐3‐phosphate dehydrogenase (gapdh)*, although the latter proved highly variable between cell-types and was discarded. Data from ZFIN, <https://zfin.org/> and from Higdon et al. (2013) quantitative profiling of melanocytes and iridophores re-expressed as digital expression.

Expression in different axial positions at 24 hpf is scored for premigratory NCCs i.e. above NT

Key: 0=no expression shown; 1=expression shown, '-' = data not available for that gene at that stage.

**Supplementary Table 2 MTE Primers**

**Supplementary Table 3 TaqMan primers**

**Supplementary Table 4 TaqMan raw data**
